# Identification of a recently dominate sublineage in *Salmonella* 4,[5],12:i:- sequence type 34 isolated from food animals in Japan

**DOI:** 10.1101/2020.10.14.338830

**Authors:** Nobuo Arai, Tsuyoshi Sekizuka, Yukino Tamamura-Andoh, Lisa Barco, Atsushi Hinenoya, Shinji Yamasaki, Taketoshi Iwata, Ayako Watanabe-Yanai, Makoto Kuroda, Masato Akiba, Masahiro Kusumoto

**Author notes:** Address for correspondence: Masahiro Kusumoto, National Institute of Animal Health, National Agriculture and Food Research Organization, 3-1-5 Kannondai, Tsukuba, Ibaraki 305-0856, Japan.

## Abstract

*Salmonella* Typhimurium and its monophasic variant (*Salmonella* 4,[5],12:i:-) are classified into nine clades. Clade 9 is composed of ST34 strains, including the *Salmonella* 4,[5],12:i:-European clone, which has rapidly disseminated in humans and animals worldwide since the 2000s. To reveal the microevolution of ST34/clade 9, we analyzed the whole-genome-based phylogenetic relationships of 230 *Salmonella* Typhimurium and *Salmonella* 4,[5],12:i:-strains isolated between 1998 and 2017 in Japan and other countries. We identified clade 9-2, a novel sublineage derived from the conventional *Salmonella* 4,[5],12:i:-clade 9 that exclusively consists of Japanese isolates. Clade 9-2 was generated by mobile genetic element-mediated stepwise genome reductions caused by IS*26* and prophages. Furthermore, the recently identified strains developed further resistance to antimicrobials via the acquisition of resistance genes. During this microevolution, *Salmonella* 4,[5],12:i:-clade 9-2 has dominated among recent isolates from food animals in Japan via the emergence of clonally expanding and/or multiple host-adapted subclades.

## Introduction

*Salmonella enterica* subsp. *enterica* serovar Typhimurium and its monophasic variant (*Salmonella* 4,[5],12:i:-) are two of the most common gastrointestinal pathogens for both humans and animals worldwide (1, 2). *Salmonella* 4,[5],12:i:-infections have been increasingly occurring in European countries since the 2000s and have since spread to the United States, South America, Australia, and Asia (3-9). A European clone is one of the best studied *Salmonella* 4,[5],12:i:-isolates because of its rapid and worldwide dissemination (6, 9-11). Swine was reported as the main reservoir of this clone, and the contaminated products are considered causative agents of food-borne infection (12). The European clone is characterized by sequence type (ST) 34 and possession of two mobile genetic elements (MGEs) on the chromosome: a composite transposon, which is responsible for resistance to core antimicrobials including ampicillin, streptomycin, sulfonamides, and tetracycline (R-type ASSuT), and an 81-kb integrative and conjugative element (ICE) containing heavy-metal tolerance genes (10). The ST34-specific ICE was first identified in *Salmonella* 4,[5],12:i:-and was named *Salmonella* genomic island 3 (SGI-3) in 2016 (10). However, Moreno Switt et al. identified nine genomic islands in several *Salmonella* serovars and designated them as SGI-2 to SGI-10 (13). Therefore, we propose to redesignate the ST34-specific ICE as ICEmST in the present study because the SGI number-based discrimination of *Salmonella* ICE is confusing.

In Japan, cases of cattle and swine salmonellosis caused by *Salmonella* 4,[5],12:i:-have increased over the past decade (14, 15). We previously classified 119 *Salmonella* Typhimurium/4,[5],12:i:-strains isolated in Japan and Italy into nine clades by whole-genome-based phylogenetic analysis (16). Among the nine clades, clade 9 was composed of *Salmonella* Typhimurium/4,[5],12:i:- and regarded as the European clone. We also developed a PCR-based clade typing method and classified another 955 *Salmonella* Typhimurium/4,[5],12:i:-strains isolated from animals in Japan (16). Clade 9 was the most predominant among the 955 strains and consisted of 210 strains: two *Salmonella* Typhimurium strains isolated in 1998 and 208 *Salmonella* 4,[5],12:i:-strains isolated between 2002 and 2017. Clade 9 strains were found in the 1990s and early 2000s but have dominated since approximately 2012; thus, the rapid dissemination of clade 9 strains in Japan may have occurred in the mid-2000s or later.

A major factor underlying the generation of the monophasic variant of *Salmonella* Typhimurium was the absence of *fljB* encoding the phase 2-H antigen (14). Other causes were the following amino acid substitutions in proteins that induced phase variation: A46T in FljA and R140L in Hin (14). In the *Salmonella* 4,[5],12:i:-European clone, the absence of *fljB* and the flanking region resulted in the expression of only one H antigen (10, 17), but how *fljB* and the flanking region were deleted from the European clone remains unclear.

The acquisition of resistance to clinically important antimicrobials is a well-studied factor regarding the clonal expansion of bacterial strains (18). Several studies have reported the acquisition of antimicrobial resistance (AMR) in addition to core antimicrobials in *Salmonella* 4,[5],12:i-ST34; the recent swine isolates acquired trimethoprim and/or gentamycin resistance via plasmids in the United Kingdom (19), and human isolates possessed a large IncHI2 plasmid (246 kb) carrying multiple AMR genes in Vietnam (11). MGEs are known to play an important role in the transmission of various genes, but the factors underlying the acquisition of such large plasmids by recently isolated ST34 strains have not been fully elucidated.

In the present study, we investigated the phylogenetic relationships of 230 *Salmonella* Typhimurium/4,[5],12:i:-clade 9 strains isolated from food animals in Japan and some other countries and analyzed the microevolution focusing on the generation of the monophasic variant. We report the emergence of a novel sublineage in *Salmonella* 4,[5],12:i:-clade 9 that has dominated in recent isolates from food animals in Japan.

## Materials and Methods

A total of 227 *Salmonella* Typhimurium/4,[5],12:i:-strains isolated in Japan and Italy (214 and 13 strains, respectively) were used in this study (Technical Appendix Table 1), and all strains have already been identified as belonging to ST34/clade 9 (16). Detailed information on the Japanese strains is shown in Technical Appendix Table 2. Among the 227 strains, the whole-genome sequences of 200 strains isolated in Japan were analyzed in this study, and those of the remaining 27 strains were reported in our previous study (16). We selected the whole-genome sequences of *Salmonella* 4,[5],12:i:-ST34, which are well supported by epidemiological information, from the National Center for Biotechnology Information database and added the data of the Australian strain TW-Stm6 and Chinese strains 81741 and WW012 (accession numbers CP019649.1, CP019442.1, and CP022168.1, respectively) for phylogenetic analysis.

Extraction of single nucleotide polymorphisms (SNPs) from the core genomes of the 230 strains was performed according to the procedures described in the Technical Appendix. To analyze the phylogenetic relationships, a maximum-likelihood (ML) tree was generated by using RAxML (20). To reveal the divergence history, we performed a temporal phylogenetic analysis of the SNPs of 214 Japanese strains by using BEAST (21) and estimated the years of emergence of distinct clades. The population structure was analyzed by hierBAPS (22) using a Bayesian clustering method.

AMR genes, plasmid replicon types, prophages, and ICEmST were identified in whole-genome sequences by the methods described in the Technical Appendix. Comparison of the genomic structures around *fljB* was performed by using Easyfig (23). The antimicrobial susceptibilities of the strains were analyzed by the disk diffusion and agar dilution methods according to recommendations provided by the Clinical and Laboratory Standards Institute (24, 25).

The detailed procedures and materials for all methods are described in the Technical Appendix. The intercellular transfer frequency of ICEmST was determined by a conjugation experiment described previously (26). We repeated the experiments three times for each donor and evaluated the data by one-way analysis of variance (ANOVA) with R (27).

## Results

### Phylogenetic overview of *Salmonella* Typhimurium/4,[5],12,i:-clade 9

A total of 911 informative SNPs were identified in the core genomes of four and 226 strains of *Salmonella* Typhimurium and 4,[5],12:i:-clade 9, respectively. The clade 9 strains were divided into two sublineages, designated clades 9-1 and 9-2, by hierBAPS (Figure 1).

**Figure 1.**
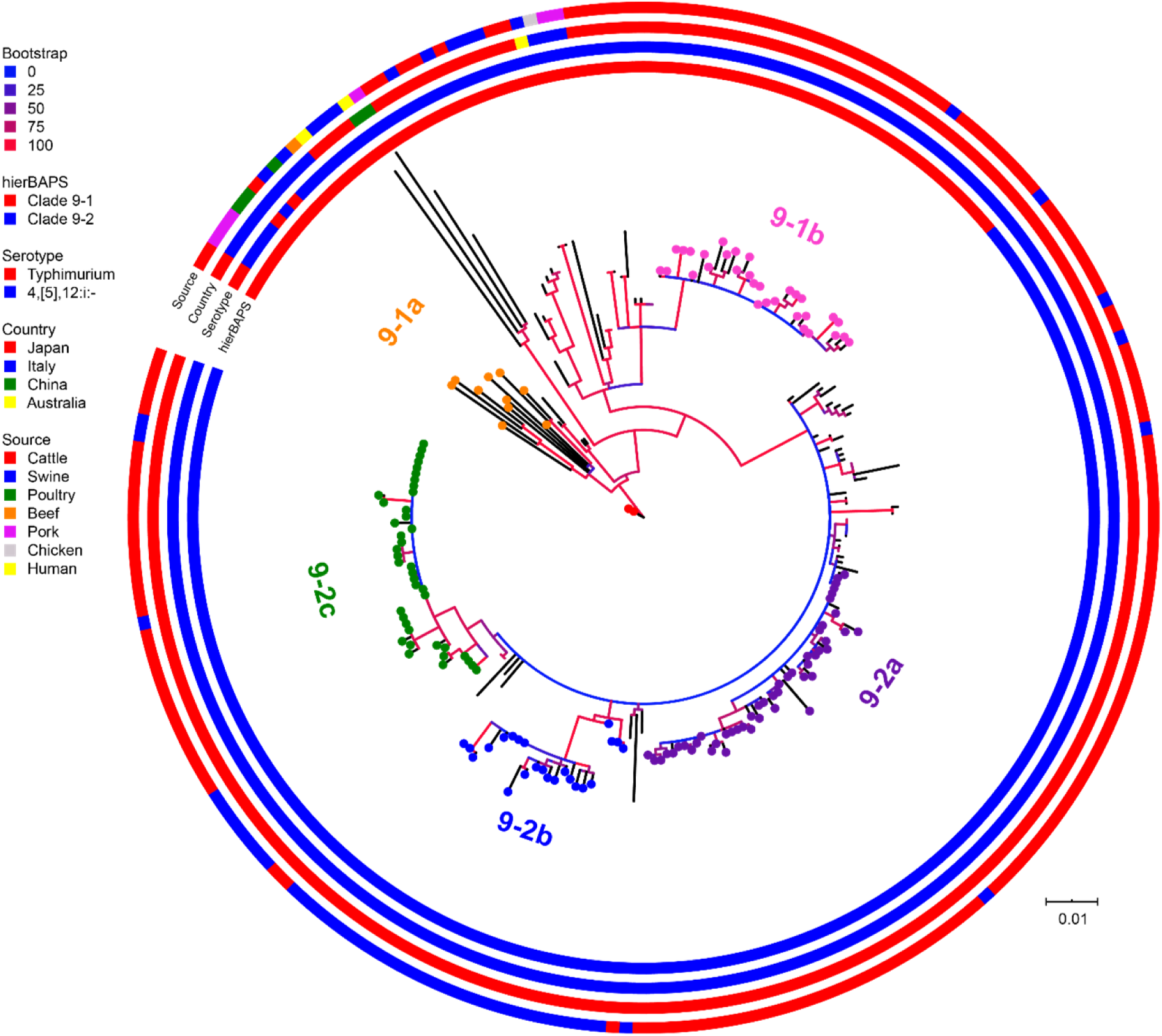
Phylogenies of *Salmonella* Typhimurium and *Salmonella* 4,[5],12:i:-ST34 isolates. A phylogenetic tree was generated using the maximum-likelihood method with 911 concatenated SNPs in the core genomic sequences of 227 wild-type strains and 3 reference strains. From the inside, the colored rings represent the hierBAPS cluster, serotype, country, and source of isolation. The red dots represent Japanese *Salmonella* Typhimurium strains, L-4126 and L-4127, isolated in 1998. The other colored dots represent strains belonging to distinctive subclades: orange, 9-1a; pink, 9-1b; purple, 9-2a; blue, 9-2b; and green, 9-2c.

Clade 9-1 was composed of four and 69 strains of *Salmonella* Typhimurium and *Salmonella* 4,[5],12:i:-, respectively, which were isolated in Japan, Italy, China, and Australia between 1998 and 2017. Clade 9-1 members were derived from the following sources: 49 from cattle/beef, 18 from swine/pork, four from poultry/chicken, and two from humans. Two *Salmonella* Typhimurium strains isolated from cattle in 1998 in Japan (Figure 1, red dots) were first branched out from the most recent common ancestor of *Salmonella* Typhimurium/4,[5],12:i:-clade 9. Italian monophasic strains were branched out and formed subclade 9-1a (Figure 1), followed by the generation of several branches consisting of the strains isolated from various sources in Japan, Italy, Australia, and China. The monophasic strains isolated from cattle (except for one from swine) in Japan between 2013 and 2017 formed a distinctive subclade, 9-1b (Figure 1).

Clade 9-2 was derived from the middle of clade 9-1 and became a predominant sublineage consisting of *Salmonella* 4,[5],12:i:-strains that were isolated from cattle and swine in Japan between 2012 and 2017. The proportion of swine strains in clade 9-2 (27%) was higher than that in subclade 9-1b (p < 0.01). Clade 9-2 included three subclades, 9-2a, 9-2b, and 9-2c, and subclades 9-2a and 9-2b were composed of the monophasic strains that were isolated from cattle (except for one from swine) and swine, respectively (Figure 1). In contrast, subclade 9-2c included strains isolated from both animals (76% in cattle, 24% in swine). Notably, clade 9-2 and subclades 9-1b, 9-2a, 9-2b, and 9-2c had high bootstrap support values in the phylogenetic tree (Figure 1).

### Temporal analysis of the emergence of *Salmonella* 4,[5],12,i:-clade 9-2 in Japan with deletion of large MGEs

Clade 9-2 was a novel sublineage derived from the conventional lineage in *Salmonella* 4,[5],12:i:-ST34 (clade 9-1) and exclusively consisted of strains isolated in Japan. To estimate the emergence times of clades 9-1 and 9-2, we analyzed a time-scaled phylogeny based on core-genome SNPs in the Japanese strains (214 strains: two *Salmonella* Typhimurium strains and 212 *Salmonella* 4,[5],12:i:-strains) by using BEAST. As shown in Figure 2, *Salmonella* 4,[5],12:i:-branched out from *Salmonella* Typhimurium (indicated with red dots) in approximately 1996. Although subclade 9-1b (Figure 2, pink dots) branched out in approximately 2009, genomic diversification of the subclade members has occurred since approximately 2012. Clade 9-2, derived from clade 9-1, was estimated to have emerged in approximately 2002. However, genomic diversification of the members of three subclades, 9-2a, 9-2b, and 9-2c (Figure 2, purple, blue, and green dots, respectively), has occurred since approximately 2012, which was approximately the same time that diversification occurred in subclade 9-1b.

**Figure 2.**
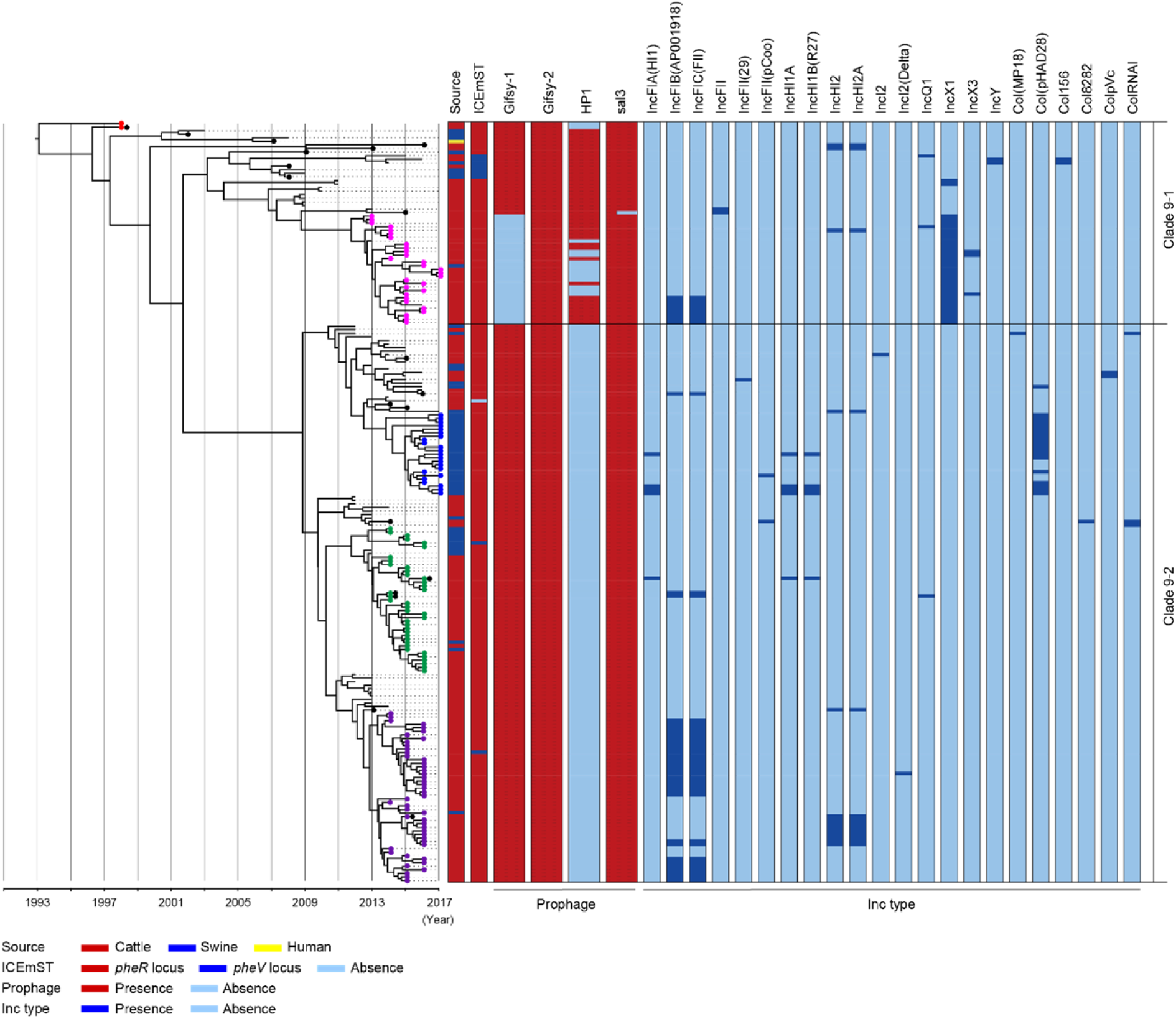
Time-scaled phylogenic analysis of *Salmonella* Typhimurium and *Salmonella* 4,[5],12:i:-strains isolated in Japan between 1998 and 2017. A phylogenetic tree was reconstructed using BEAST v1.8.2 based on 911 concatenated SNPs in the core genomic sequences of 214 wild-type strains isolated in Japan. Detailed information for each strain is described in Technical Appendix Table 2. The x-axis indicates the time of emergence for each branch. The red, pink, blue, green, and purple dots represent the strains corresponding to Figure 1. The black dots represent the strains for which the complete sequences were determined.

The prophage and plasmid replicon type distributions are also shown in Figure 2. Among 19 strains for which complete genomic sequences were determined (Figure 2, black dots), four kinds of intact prophages were identified by using PHASTER. Each of the four prophages was completely conserved in the positive strains except for the 3’-terminus of the Gifsy-1 prophage, which was deleted in only one strain (Technical Appendix Figure 1). For the other 195 strains, possession of the four prophages was determined by the detection of three genes, which were selected from the 5’-, mid-, and 3’-regions of the draft genomic sequences of each prophage. Plasmids were also detected from the draft genomic sequences as Inc types. As shown in Figure 2, subclade 9-1b lacked Gifsy-1, a common prophage in *Salmonella* Typhimurium (28), instead of possessing the IncX1 plasmid; in contrast, clade 9-2 lacked the HP1 prophage.

ICEmST, an ST34/clade 9-specific MGE, was identified in most Japanese strains at the *pheR* locus on the chromosome (Figure 2). Seven strains involved in the same branch of clade 9-1 carried ICEmST at the *pheV* locus. In clade 9-2, ICEmST was located on the *pheV* locus in two strains associated with subclades 9-2a and 9-2c. At least three intracellular transpositions of ICEmST from *pheR* to *pheV* occurred independently, and excision of ICEmST from the chromosome was observed in one strain located near subclade 9-2b on the phylogenetic tree (Figure 2). Notably, there were no significant differences in the intercellular transfer frequencies of ICEmST among the strains representing clades 9-1 and 9-2 (Technical Appendix Table 3).

### Small MGE-mediated stepwise deletion of genomic regions related to phase variation

The *fljAB* operon is replaced by a composite transposon containing two copies of IS*26* and AMR genes in *Salmonella* 4,[5],12:i:-(17), and we previously showed that this transposon is specific for *Salmonella* 4,[5],12:i:-clade 9 (16). Although the LT2 strain (*Salmonella* Typhimurium clade 3) carried only the *fljAB* operon, the L-4126 strain (*Salmonella* Typhimurium clade 9-1) isolated in 1998 in Japan was found to carry both the *fljAB* operon and the clade 9-specific transposon inserted between the *hin* and *iroB* genes (Figure 3a). The TW-Stm6 strain (*Salmonella* 4,[5],12:i:-clade 9-1) isolated in 2014 in Australia (29) carried the transposon instead of the region between the STM2760 and *hin* genes; the same region was deleted in 46 of 57 strains in clade 9-1 (Figure 3b). Additionally, the deleted region has expanded to the STM2752 gene in the L-4445 strain (*Salmonella* 4,[5],12:i:-clade 9-2) isolated in 2014 in Japan (Figure 3a). All members of clade 9-2 lacked the region between the STM2752 and *hin* genes (Figure 3b). Among all strains in clades 9-1 and 9-2, the terminal IS*26* in the clade 9-specific transposon remained intact.

**Figure 3.**
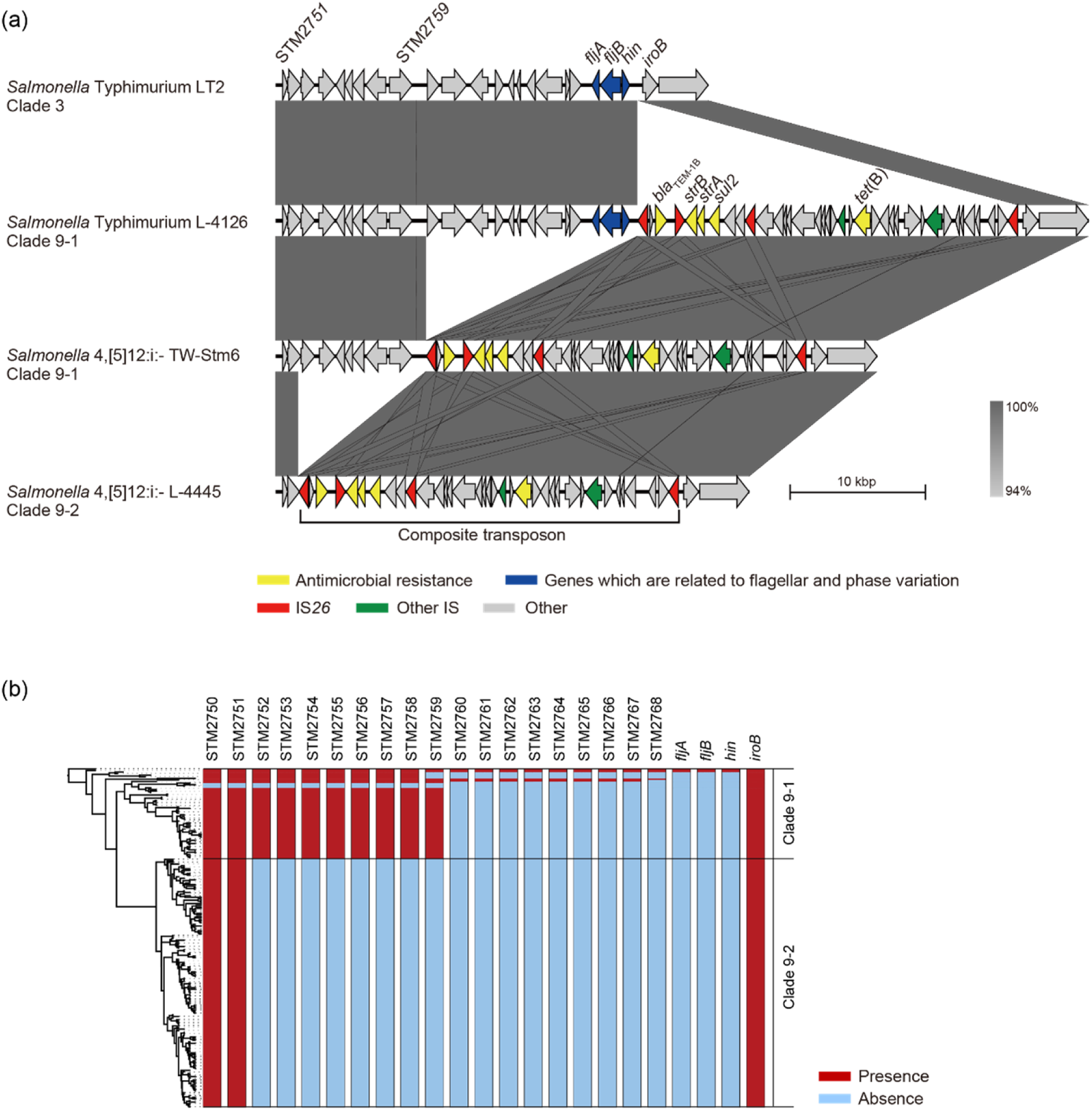
Comparison of the flanking regions of *fljB*. (a) Comparison of the genetic structures among non-clade 9 *Salmonella* Typhimurium, clade 9-1 *Salmonella* Typhimurium, clade 9-1 *Salmonella* 4,[5],12:i:-, and clade 9-2 *Salmonella* 4,[5],12:i:-. (b) Gene repertoires located between the STM2750 and *iroB* genes are shown beside the phylogenetic tree represented in Figure 2.

### Acquisition of AMR in clade 9 strains

The five AMR genes, *bla*_TEM-1B_, *strA, strB, sul2*, and *tet*(B), carried on the clade 9-specific transposon were detected in all 214 strains in clades 9-1 and 9-2. However, some genes were broken, and 185 strains (86%) showed ASSuT (Figure 4). In addition to the five genes, *dfrA12, floR*, and *cmlA1* were detected at relatively high levels (25%, 22%, and 21%, respectively). These resistance genes for trimethoprim and phenicols were found in large plasmids (129 to 230 kb in size), which belong to replicon types IncFI and IncHI and carry multiple AMR genes identified in the complete genome-determined strains (Technical Appendix Table 4). The strains showing resistance to chloramphenicol and trimethoprim in addition to ASSuT (R-type ASSuTCTm) accounted for 18% of the total, and the rate of this R-type has been rapidly increasing (2.6% before 2014 and 26% after 2015) (Figure 4). The third largest R-type was ASSuTCTmK, which represented the strains showing resistance to kanamycin in addition to ASSuTCTm. The strains showing both ASSuTCTm and ASSuTCTmK possessed plasmids ranging from <40 to 208 kb (Technical Appendix Table 2).

**Figure 4.**
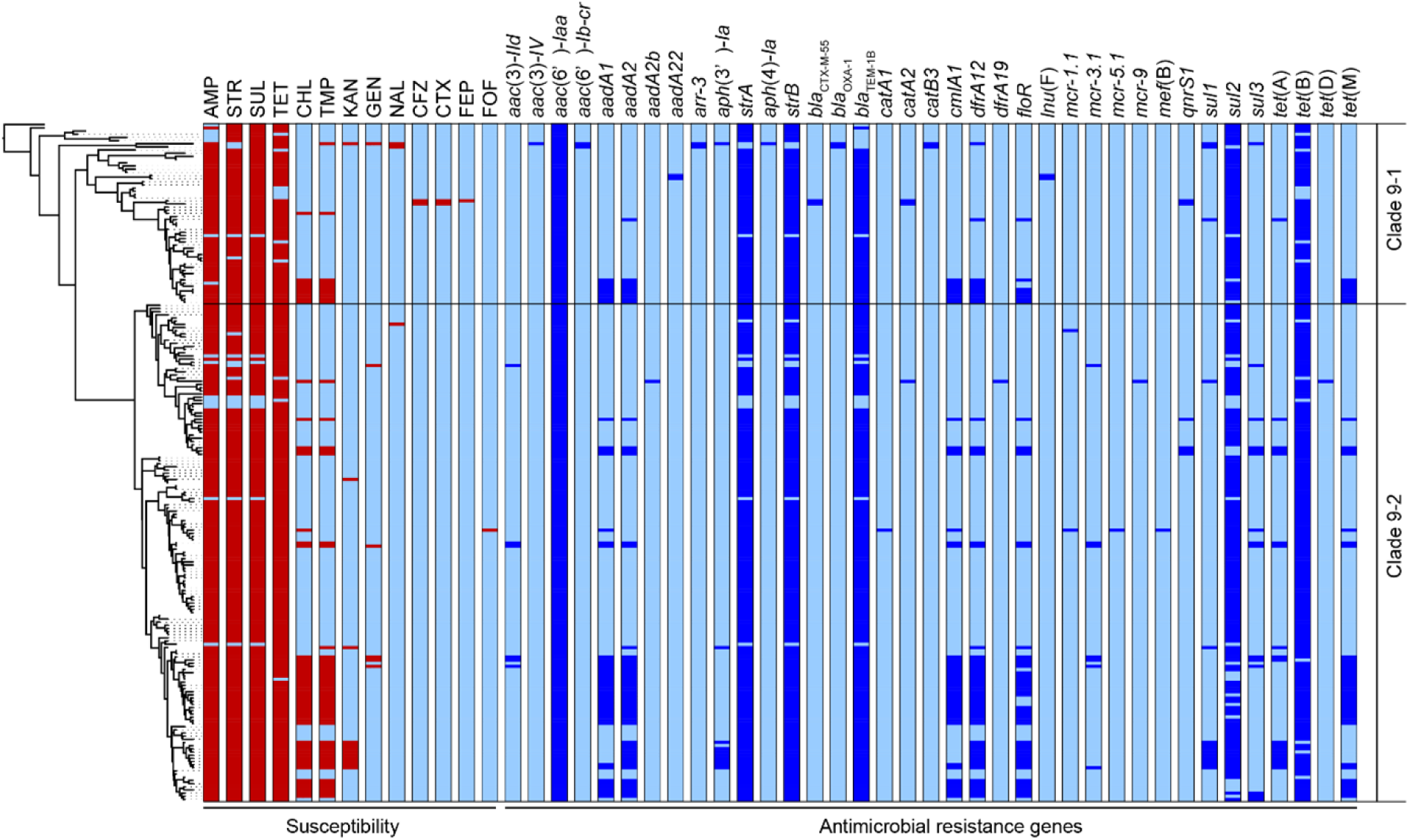
The antimicrobial susceptibilities and distributions of antimicrobial resistance genes are shown beside the phylogenetic tree represented in Figure 2. The red and light blue colors relating to susceptibility (left side) indicate the resistance and susceptibility to each antimicrobial, respectively. The blue and light blue colors relating to the antimicrobial resistance genes (right side) indicate the presence and absence of each gene, respectively.

Importantly, an extended-spectrum β-lactamase (ESBL) gene, *bla*CTX-M-55, was detected in two strains belonging to clade 9-1 (L-4071 and L-4526), which were isolated from cattle in Japan, and both strains showed resistance to cefazolin and cefotaxime (Figure 4). Furthermore, the L-4071 strain exhibited resistance to cefepime and carried *bla*CTX-M-55 on the chromosome, and IS*26* and the series of genes associated with conjugal transfer were located around *bla*CTX-M-55. In addition, the sequence of the integrated region showed high similarity to the *E. coli* plasmid pHNHN21 (accession number KX24667.1).

Colistin resistance genes were detected in only clade 9-2. Totals of two, seven, one, and one strains possessed *mcr-1, mcr-3, mcr-5*, and *mcr-9*, respectively (Figure 4). All but the L-4795 strain, which possessed *mcr-9* and was isolated from swine, were isolated from cattle. As shown in Technical Appendix Table 4, *mcr*-positive plasmids, pSAL4445-1, pSAL4567-1, pSAL4596-1, and pSAL4605-1, were identified in the complete genome-determined strains. In the four plasmids, pSAL4596-1 with replicon types IncFIA(HI1), IncHI1A, and IncHI1B(R27) carried both *mcr-1* and *mcr-5*. The other three plasmids with replicon types IncFIB(AP001918) and IncFIC(FII) carried *mcr-3*.

## Discussion

*Salmonella* 4,[5],12:i:-ST34/clade 9 was first isolated in approximately 2005 in Europe and spread across European countries (3-5, 10), followed by North and South America, Asia, and Australia (6-9). In the present study, we elucidated the phylogenetic relationship of *Salmonella* Typhimurium clade 9 and its monophasic variant caused by MGE-mediated microevolution and identified the emergence of a recently dominated sublineage, designated clade 9-2, in conventional clade 9 of *Salmonella* 4,[5],12:i:-(clade 9-1) in Japan. Clade 9-1 included the *Salmonella* 4,[5],12:i:-European clones and diverse strains derived from various sources and countries (Figure 1). *Salmonella* 4,[5],12:i:-was derived from *Salmonella* Typhimurium in approximately 1996, and the clade 9-2 was estimated to have emerged in approximately 2002 (Figure 2). Therefore, clade 9-2 might have already been generated but had not appeared as an isolate before the isolation of the first European clone. In previous reports about phylogenies in *Salmonella* Typhimurium (10) and *Escherichia coli* (30), clusters with a maximum node-to-tip distance of up to 70 SNPs were considered clonally expanding clades. Clades 9-1 and 9-2 included the epidemic subclades 9-1b, 9-2a, 9-2b, and 9-2c in the range of 63, 67, 56, and 47 SNPs, respectively, suggesting that each subclade is associated with independent clonal expansion. Interestingly, genomic diversification in the four subclades has occurred around the same time since approximately 2012 (Figure 2). In Japan, the incidence of salmonellosis in food animals caused by *Salmonella* 4,[5],12:i:-clade 9 has also increased since 2012 (16). The clonally expanding four subclades were suggested to have diversified via the dissemination of *Salmonella* 4,[5],12:i:-clade 9 in Japan but did not appear immediately and were isolated mostly between 2015 and 2017.

Although swine was identified as the major reservoir of *Salmonella* 4,[5],12:i:-ST34/clade 9 in Europe (19, 31), clade 9 strains have been isolated mainly from cattle in Japan (16). As shown in Figure 2, clade 9-2 and the clonally expanding subclades 9-1b, 9-2a, 9-2b, and 9-2c were derived from branches of cattle isolates. Most strains belonging to subclades 9-1b and 9-2a were isolated from cattle. In contrast, all and 24% of subclades 9-2b and 9-2c, respectively, consisted of swine isolates. These observations suggested that clade 9-2 has acquired the ability to adapt effectively into swine in addition to cattle.

The prophage repertoire was noted as a remarkable difference between sporadic strains and the clonally expanding subclades, as subclade 9-1b and clade 9-2 lacked the Gifsy-1 and HP1 prophages from the chromosome, respectively (Figure 2). Alternatively, the clonally expanding subclades involved multiple plasmids, such as replicon types IncFI, IncHI, IncXI, and Col (Figure 2). In a study on *E. coli*, genome reduction led to highly efficient plasmid acquisition and accurate propagation of foreign genes (32). Therefore, the deletion of the prophages may have contributed to the clonal expansion by the acquisition of some genes via MGEs that promoted subclade dissemination. As reported by Petrovska et al., the *sopE* gene was identified in the mTmV prophage and considered to be a virulence gene in *Salmonella* 4,[5],12:i:-(10). In our data, three and one strains associated with clades 9-1 and 9-2 possessed the *sopE* gene, respectively, but they were neither located on the mTmV prophage nor found in the clonally expanding subclades.

*Salmonella* 4,[5],12:i:-is considered a monophasic variant of *Salmonella* Typhimurium because of its close genetic relationship (16); however, little is known about how the monophasic variant was generated. As shown in Figure 3, stepwise deletions of the genomic regions related to the phase variation were observed in *Salmonella* Typhimurium clade 9-1 and *Salmonella* 4,[5],12:i:-clades 9-1 and 9-2. The STM2759 and STM2751 genes were adjacent to IS*26*, which was located at the end of the clade 9-specific transposon, in the TW-Stm6 and L-4445 strains, respectively. IS elements are the simplest MGEs and are known to induce a variety of genomic rearrangements, such as deletions, inversions, and duplications. Recently, He et al. demonstrated DNA deletions caused by the intramolecular transposition of IS*26* that consisted of cleavage at IS-ends by transposase, generation of 3’-OH groups to attack the target site on the same strand, circularization of a region between IS and the target site, and removal of the region from original DNA (33). In our study, there was no IS element around the *fljAB* operon except for IS*26* at the ends of the transposon (Figure 3), suggesting that the stepwise deletions were caused not by homologous recombination but rather by intramolecular transposition of IS*26*. As a result of these microevolutions, *Salmonella* 4,[5],12:i:-has lost genes encoding a phosphoenolpyruvate-dependent sugar phosphotransferase (PTS) system and some hexulose synthases in clade 9-2. Further research will be required to clarify the effect of these genes on the phenotype of *Salmonella enterica*.

Although *Salmonella* 4,[5],12:i:-ST34 typically shows ASSuT (10, 17), several studies have highlighted the emergence of trimethoprim-resistant European clones in food animals (19, 34). Our data also showed that ASSuTCTm was the second largest R-type in *Salmonella* 4,[5],12:i:-ST34/clade 9 and that it has been increasingly isolated in recent years. This observation is consistent with the fact that phenicols and trimethoprim have been used for food animals at a certain level in Japan (35). In addition, the *bla*CTX-M-55 gene was detected in two strains that were resistant to third-generation cephalosporins (cefotaxime) and were closely related to subclade 9-1b on the phylogenetic tree (Figures 2 and 4). One of the two strains carried *bla*CTX-M-55 on the chromosome and showed further resistance to fourth-generation cephalosporins (cefepime). During the past decade, *bla*CTX-M-55 has been increasingly found in *Enterobacteriaceae* isolated from humans, animals, and the environment (36). Recently, *mcr* genes were found in *Salmonella* Typhimurium/4,[5],12:i:-ST34 isolated from humans and swine in North America, Europe, Australia, and Asian countries other than Japan (8, 11, 34, 37, 38). The present study is the first to identify *mcr-1, mcr-3, mcr-5*, and/or *mcr-9* in *Salmonella* 4,[5],12:i:-ST34 disseminated into food animals in Japan. The *mcr* genes were identified in only clade 9-2 strains isolated after 2014, suggesting that acquisition of the plasmid-encoded *mcr* genes has occurred during the process of microevolution involving genome reduction. Obviously, third- and fourth-generation cephalosporins and colistin are important antibiotics for human health. The progression of AMR in *Salmonella* 4,[5],12:i:-ST34/clade 9 should be monitored continuously as a public health concern.

In conclusion, we identified a novel sublineage, clade 9-2, in *Salmonella* 4,[5],12:i:-ST34/clade 9 caused by several MGE-mediated microevolution steps (Figure 5). First, *Salmonella* Typhimurium clade 9-1 was derived from the ancestral clone by acquiring ICEmST and the clade 9-specific transposon on the chromosome. Second, *Salmonella* Typhimurium clade 9-1 lost the *fljAB* operon by the intramolecular transposition of IS*26* and converted to the monophasic variant *Salmonella* 4,[5],12:i:-clade 9-1. Finally, *Salmonella* 4,[5],12:i:-clade 9-2 emerged from clade 9-1 via additional genome reductions caused by the IS*26*-mediated deletion adjacent to the transposon and loss of the HP1 prophage. Furthermore, *Salmonella* 4,[5],12:i:-subclade 9-1b reduced the genome size via loss of the Gifsy-1 prophage before clonal expansion. Alternatively, some strains have developed more resistance to antimicrobials through the acquisition of plasmids and resistance genes. During this microevolution, *Salmonella* 4,[5],12:i:-clade 9-2 has dominated in recent isolates from food animals in Japan via the emergence of clonally expanding and/or multiple host-adapted subclades. Recently, strains that were considered monophasic variants of known serovars other than Typhimurium have been isolated from humans and animals. In these strains, MGE-mediated microevolution involving genome reduction followed by acquisition of effective genes may be of the same importance as that shown in our study.

**Figure 5.**
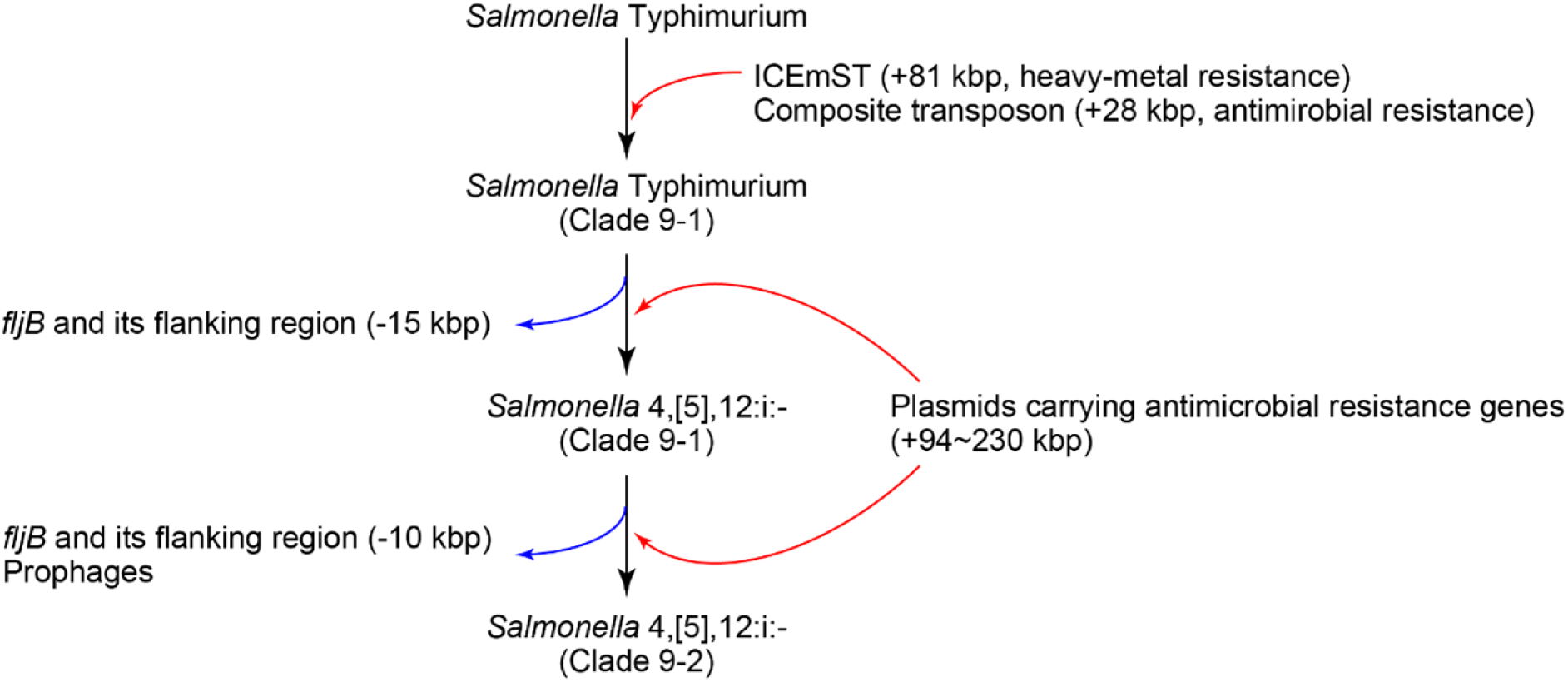
Schematic overview of the microevolution of epidemic lineages of *Salmonella* Typhimurium and *Salmonella* 4,[5]12:i:-. The black, red, and blue arrows indicate the flow of microevolution, acquisition of foreign genes, and genome reduction, respectively.

## Acknowledgment

We are grateful to the prefectural livestock hygiene service centers for providing *Salmonella* isolates.

This work was supported by KAKENHI grant number JP19J10415 provided by the Japan Society for the Promotion of Science (JSPS). This study was also supported by a grant-in-aid for the Research Program on Emerging and Reemerging Infectious Diseases (15fk0108021h0002) provided by the Japan Agency for Medical Research and Development (AMED).

## Biography

Dr. Arai is a research scientist at the National Institute of Animal Health, National Agriculture and Food Research Organization in Ibaraki, Japan. His main research interest is molecular epidemiological studies using high-throughput sequencing methods to understand the dissemination of enteropathogenic bacteria among humans and food animals.

