## Supplemental Tables and Figures. for "Identification of a recently dominate sublineage in *Salmonella* 4,[5],12:i:- sequence type 34 isolated from food animals in Japan"

**Technical Appendix**

**Whole-genome sequencing and genome assembly.** Genomic DNA extraction of each isolate was performed as described in our previous study (1). The sequencing library was prepared using a Nextera XT DNA sample preparation kit (Illumina, Inc., San Diego, CA), and sequencing was performed using an Illumina NextSeq 500 sequencer (Illumina) according to the manufacturer's instructions. The de novo assembly of all short reads was performed using A5 MiSeq program software version 20140604 (2).

To determine the complete genomic sequences of one and 18 isolates of *Salmonella* Typhimurium and *Salmonella* 4,[5],12:i:-, respectively, the high-molecular weight DNA was extracted followed by long-read sequencing using a PacBio Sequel sequencer (Sequel SMRT Cell 1M v2 [four/tray]; Sequel Sequencing Kit v2.1; insert size, approximately 10 kb). The long-read libraries were prepared with a SMRTbell library using a SMRTbell Template Prep Kit 1.0 (PacBio, Menlo Park, CA) with barcoding adapters according to the manufacturer's instructions. These sequencing data were produced with more than 50-fold coverage, and preassembled reads were generated using SMRT Link software v5. De novo assembly with preassembled reads was performed using Canu version 1.4 [PMID: 28298431], Minimap version 0.2-r124 [PMID: 27153593], racon version 1.1.0 [PMID: 28100585], and Circlator version 1.5.3 [PMID: 26714481]. Error correction of assembled sequences was performed using Pilon version 1.18 [PMID: 25409509] with Illumina short reads previously deposited in the Sequence Read Archive (SRA) database (3).

***In silico* multilocus sequence typing (MLST).** MLST was performed using the nucleotide sequences of seven housekeeping genes, *aroC*, *dnaN*, *hemD*, *hisD*, *purE*, *sucA*, and *thrA*, found on draft genome contigs, according to protocols available in the MLST database (http://mlst.warwick.ac.uk/mlst/dbs/Senterica).

**SNP detection and phylogenetic analysis.** The paired-end Illumina sequence data were mapped onto a complete genomic sequence of the L-3837 strain using the bwa-mem program (4). SNPs were extracted using VarScan v2.3.4 (5). Repeat and prophage regions were searched by the NUCmer (MUMmer 3.0) and PHAST (6) programs, respectively, followed by exclusion of SNPs in these regions. In all of the 230 tested isolates, 911 SNP sites were identified. RAxML (7) was used to reconstruct a maximum-likelihood phylogenetic tree based on the concatenated 911 core-genome SNPs. The general time reversible (GTR)-GAMMA model was used as a nucleotide substitution model with 1,000 bootstraps. The phylogenetic tree was visualized and annotated by using iTOL (8). The population structure was analyzed with hierBAPS software (9) using a Bayesian clustering method.

**Temporal Bayesian phylogenetic analysis.** We performed temporal analysis of 214 clade 9 strains isolated in Japan by using BEAST v1.8.2 (10). The GTR model of nucleotide substitution was selected as the best fit model under the Akaike information criterion (AIC) using jmodeltest v2.1.10 (11, 12). We compared 32 combinations of four types of clock models and eight types of tree priors. The isolation year was used to calibrate the time scale of the tree. Markov chain Monte Carlo (MCMC) analysis of each combination was run with a $10^8$ chain length, sampling each 1,000 steps to ensure convergence. Model performance was assessed by Tracer v1.6.0 (http://tree.bio.ed.ac.uk/software/tracer/), which indicated adequate convergence of run statistics and all effective sample size values greater than 200. As a result, the combination was determined to be an exponential relaxed clock model and a coalscent: exponential growth (tree prior). Triplicate runs were performed, and each lineage was combined with LogCombiner v1.8.0 (implemented in BEAST) with the first 10% of the stats in each chain discounted as a burn in. The trees produced by BEAST were summarized by a single maximum clade credibility tree using TreeAnnotator v1.8.0 (implemented in BEAST), followed by visualization with FigTree v1.4.2 (http://tree.bio.ed.ac.uk/software/figtree/).

**Detection of antimicrobial resistance genes, plasmid Inc type, phages, and ICEmST.** Antimicrobial resistance genes were searched by ResFinder (13) with the following thresholds: minimum coverage length ≥80% and nucleotide sequence identity ≥90%.

Plasmid Inc types were searched by using PlasmidFinder (14) with the following thresholds: minimum coverage length ≥80% and nucleotide sequence identity ≥90%.

Predicted prophage sequences were detected by PHASTER (15), and the regions that were detected as intact by PHASTER were identified as prophages. Three ORFs from each prophage were used for homology searches by the NUCmer alignment program from MUMmer in GENETYX version 13 (GENETYX Co., Ltd., Tokyo, Japan). The following ORFs were used as query sequences (the positions in each prophage are indicated as yellow arrows in Technical Appendix Figure 1): Gifsy-1 in the L-4126 strain (accession number AP023291), SAL4126_11860, ASL4126_12080, and SAL4126_12320; Gifsy-2 in the L-4126 strain, SAL4126_27940, SAL4126_28160, and SAL4126_28280; HP1 in the L-4526 strain (accession number AP023303), SAL4526_06240, SAL4526_06470, and SAL4526_06570; sal3 in the L-4126 strain, SAL4126_18170, SAL4126_18520, and SAL4126_18590.

A homology search was performed using the NUCmer alignment program to detect ICEmST in the *pheR* and *pheV* loci on the chromosome. The following regions were used as query sequences: the flanking region of ICEmST at the *pheR* locus (accession number AP019375.1), nucleotides (nt) 4,336,860 to 4,337,765 and 4,417,770 to 4,418,724; and the flanking region of ICEmST at the *pheV* locus (accession number AP019374.1), nt 800,391 to 801,295 and 881,300 to 882,199. The detection thresholds were as follows: minimum coverage length ≥90% and nucleotide sequence identity ≥90%.

**Antimicrobial susceptibility testing.** The Kirby-Bauer disk diffusion test was performed using a Muller-Hinton agar plate (Becton, Dickinson and Company, Franklin Lakes, NJ) according to the Clinical and Laboratory Standards Institute (CLSI) standards (16) using the following 16 antimicrobials: ampicillin (10 µg), streptomycin (10 µg), tetracycline (30 µg), chloramphenicol (30 µg), sulfamethoxazole-trimethoprim (23.75/1.25 µg), kanamycin (30 µg), gentamicin (10 µg), nalidixic acid (30 µg), ciprofloxacin (5 µg), cefazolin (30 µg), cefotaxime (30 µg), cefepime (30 µg), cefoxitin (30 µg), fosfomycin (50 µg), imipenem (10 µg) and meropenem (10 µg). The MIC of sulfamethoxazole (Fujifilm Wako Pure Chemical Co., Osaka, Japan) was determined by the agar dilution method (17). Overnight cultures of each strain in LB broth were diluted to approximately $10^6$ cfu/ml, and 5-µl portions were spotted onto Mueller-Hinton agar plates (Becton Dickinson) supplemented with sulfamethoxazole (4, 8, 16, 32, 64, 128, 256, and 512 µg/mL). The spotted plates were incubated at 35°C for 20 hours. The *Escherichia coli* ATCC25922 strain was used for quality control. According to the CLSI criterion (18), strains with an MIC ≥512 µg/mL were considered sulfamethoxazole resistant.

**Plasmid profile.** Plasmids were extracted from *Salmonella* strains by the alkaline lysis method as previously described (19). Plasmid DNA was separated by 0.8% agarose gel electrophoresis in 1x Tris Acetic acid EDTA buffer, pH 8.5 at 100 V for 35 minutes. Plasmid sizes were determined by comparison with plasmids of known sizes as follows: pMAK1; 208 kb (accession number AB366440.1), pMAK2; 62 kb (accession number AB366441.1), pMAK3; 40 kb (accession number AB366442.1), pSLT; 94 kb (accession number AE006471.2), and pSN; 176 kb.

**Homology search for genes in the *fljB* flanking region and pairwise alignment of the DNA sequences.** Nucleotide sequences of the genes from STM2750 to STM2773 in the *Salmonella* Typhimurium LT2 strain were used as query sequences. The detection thresholds were as follows: minimum coverage length ≥80% and nucleotide sequence identity ≥85%. Pairwise alignment of the *fljB* flanking region was performed using the BLAST search tool, followed by a homology search using the NUCmer alignment program from MUMmer in GENETYX version 13. The alignment image was visualized by using Easyfig version 2.2.2 (20).

16. Clinical Laboratory Standards Institute. Performance standerds for antimicrobial disk susceptibility tests; approved standard, 11th ed, document M02-A11. Wayne, PA. 2012.

17. Clinical Laboratory Standards Institute. Performance standards for antimicrobial susceptibility testing: 24th informational supplement. Document M100–S24. Wayne, PA. 2014.

18. Clinical Laboratory Standards Institute. CLSI M100-ED30:2020 Performance Standards for Antimicrobial Susceptibility Testing, 30th Edition, Wayne, PA. 2020.

19. Kado CI, Liu ST. Rapid procedure for detection and isolation of large and small plasmids. J Bacteriol. 1981;145:1365-73.

20. Sullivan MJ, Petty NK, Beatson SA. Easyfig: a genome comparison visualizer. Bioinformatics. 2011;27:1009-10.

**Technical Appendix Table 1.** Origin, year of isolation, number of strains of *Salmonella* Typhimurium and *Salmonella* 4,[5],12:i:- used in this study

| Serotype | Country | Origin | Year of isolation | No. of isolates |
|---|---|---|---|---|
| Typhimurium | Japan | Cattle | 1998 | 2 |
| | | Subtotal | | 2 |
| | Italy | Poultry | 2011 | 1 |
| | | Swine | 2012 | 1 |
| | | Subtotal | | 2 |
| 4,[5],12:i:- | Japan | Cattle | 2008-2017 | 160 |
| | | Swine | 2002-2017 | 51 |
| | | Human | 2007 | 1 |
| | | Subtotal | | 212 |
| | Italy | Cattle | 2011 | 1 |
| | | Swine | 2011 | 1 |
| | | Poultry | 2010-2012 | 4 |
| | | Meats | 2011 | 6 |
| | | Subtotal | | 11 |
| | | Total | | 227 |

**Technical Appendix Table 2.** Characteristics of *Salmonella* Typhimurium and *Salmonella* 4,[5],12:i:- Japanese strains used for whole genome sequencing

| Strain | Serotype | Clade | Origin | Year | Sequence Type | ICEmST (locus)* | Plasmid profile † | Antimicrobial resistance pattern ‡ | BioSample/Experiment§ accession no. or reference |
|---|---|---|---|---|---|---|---|---|---|
| L-4127 | Typhimurium | 9-1 | Cattle | 1998 | 34 | + (*pheR*) | ND | STR, SUL, TET | 19 |
| L-4126 | Typhimurium | 9-1 | Cattle | 1998 | 34 | + (*pheR*) | ND | AMP, STR, SUL, TET | 19 |
| L-3845 | 4,[5],12:i:- | 9-1 | Swine | 2003 | 34 | + (*pheR*) | ND | STR, SUL, TET | SAMD00234571/DRX226576 |
| L-3844 | 4,[5],12:i:- | 9-1 | Swine | 2002 | 34 | + (*pheR*) | ND | STR, SUL | 19 |
| L-3846 | 4,[5],12:i:- | 9-1 | Swine | 2008 | 34 | + (*pheR*) | ND | STR, SUL, TET | 19 |
| L-3835 | 4,[5],12:i:- | 9-1 | Huma | 2007 | 34 | + (*pheR*) | ND | STR, SUL, TET | 19 |
| L-4681 | 4,[5],12:i:- | 9-1 | Cattle | 2016 | 34 | + (*pheR*) | 176 kb | AMP, SUL, TET, TMP, KAN, GEN, NAL | SAMD00234738/DRX226743 |
| L-4551 | 4,[5],12:i:- | 9-1 | Cattle | 2013 | 34 | + (*pheR*) | 176 kb | AMP, SUL, TET, NAL | SAMD00234630/DRX226635 |
| L-3841 | 4,[5],12:i:- | 9-1 | Swine | 2009 | 34 | + (*pheR*) | ND | AMP, STR, SUL | 19 |
| L-4678 | 4,[5],12:i:- | 9-1 | Cattle | 2015 | 34 | + (*pheV*) | ND | AMP, STR, SUL, TET | SAMD00234736/DRX226741 |
| L-4686 | 4,[5],12:i:- | 9-1 | Cattle | 2016 | 34 | + (*pheV*) | 94 kb, <40 kb (2) | AMP, STR, SUL, TET | SAMD00234742/DRX226747 |
| L-4684 | 4,[5],12:i:- | 9-1 | Swine | 2014 | 34 | + (*pheV*) | 176 kb | AMP, STR, SUL, TET | SAMD00234740/DRX226745 |
| L-3837 | 4,[5],12:i:- | 9-1 | Cattle | 2008 | 34 | + (*pheV*) | ND | AMP, STR, SUL, TET | 19 |
| L-3802 | 4,[5],12:i:- | 9-1 | Swine | 2009 | 34 | + (*pheV*) | ND | AMP, STR, SUL, TET | SAMD00234570/DRX226575 |
| L-3801 | 4,[5],12:i:- | 9-1 | Swine | 2009 | 34 | + (*pheV*) | ND | AMP, STR, SUL, TET | SAMD00234569/DRX226574 |
| L-3838 | 4,[5],12:i:- | 9-1 | Swine | 2008 | 34 | + (*pheV*) | ND | AMP, STR, SUL, TET | 19 |
| L-4439 | 4,[5],12:i:- | 9-1 | Cattle | 2011 | 34 | + (*pheR*) | 62 kb | AMP, STR, SUL, TET | SAMD00234597/DRX226602 |
| L-4547 | 4,[5],12:i:- | 9-1 | Cattle | 2011 | 34 | + (*pheR*) | 40 kb | AMP, STR, SUL, TET | SAMD00234626/DRX226631 |
| L-4546 | 4,[5],12:i:- | 9-1 | Cattle | 2010 | 34 | + (*pheR*) | ND | AMP, STR, SUL, TET | SAMD00234625/DRX226630 |
| L-4438 | 4,[5],12:i:- | 9-1 | Cattle | 2010 | 34 | + (*pheR*) | ND | AMP, STR, SUL, TET | SAMD00234596/DRX226601 |
| L-4825 | 4,[5],12:i:- | 9-1 | Cattle | 2009 | 34 | + (*pheR*) | <40 kb (2) | AMP, STR, SUL | SAMD00234766/DRX226771 |
| L-4824 | 4,[5],12:i:- | 9-1 | Cattle | 2009 | 34 | + (*pheR*) | <40 kb (2) | AMP, STR, SUL | SAMD00234765/DRX226770 |
| L-4827 | 4,[5],12:i:- | 9-1 | Cattle | 2009 | 34 | + (*pheR*) | <40 kb (2) | AMP, STR, SUL | SAMD00234768/DRX226773 |
| L-4826 | 4,[5],12:i:- | 9-1 | Cattle | 2009 | 34 | + (*pheR*) | <40 kb (2) | AMP, STR, SUL | SAMD00234767/DRX226772 |
| L-4071 | 4,[5],12:i:- | 9-1 | Cattle | 2013 | 34 | + (*pheR*) | ND | AMP, STR, SUL, TET, CFZ, CTX, FEP | 19 |
| L-4526 | 4,[5],12:i:- | 9-1 | Cattle | 2015 | 34 | + (*pheR*) | ND | AMP, STR, SUL, TET, CFZ, CTX | SAMD00234616/DRX226621 |
| L-4611 | 4,[5],12:i:- | 9-1 | Cattle | 2013 | 34 | + (*pheR*) | 40 kb | AMP, STR, SUL, TET | SAMD00234681/DRX226686 |

*Continued on following pages*

**Technical Appendix Table 2**-Continued

| Strain | Serotype | Clade | Origin | Year | Sequence Type | ICEmST (locus)* | Plasmid profile† | Antimicrobial resistance pattern‡ | BioSample/Experiment§ accession no. or reference |
|---|---|---|---|---|---|---|---|---|---|
| L-4613 | 4,[5],12:i:- | 9-1 | Cattle | 2013 | 34 | + (*pheR*) | 40 kb | AMP, STR, SUL, TET | SAMD00234683/DRX226688 |
| L-4556 | 4,[5],12:i:- | 9-1 | Cattle | 2014 | 34 | + (*pheR*) | 40 kb | AMP, STR, SUL, TET, CHL, TMP | SAMD00234634/DRX226639 |
| L-4563 | 4,[5],12:i:- | 9-1 | Cattle | 2014 | 34 | + (*pheR*) | 40 kb, <40 kb (2) | AMP, STR, SUL, TET | SAMD00234641/DRX226646 |
| L-4566 | 4,[5],12:i:- | 9-1 | Cattle | 2013 | 34 | + (*pheR*) | 176 kb, 40 kb | AMP, STR, SUL, TET | SAMD00234644/DRX226649 |
| L-4444 | 4,[5],12:i:- | 9-1 | Cattle | 2014 | 34 | + (*pheR*) | 40 kb | AMP, STR, SUL, TET | SAMD00234602/DRX226607 |
| L-4562 | 4,[5],12:i:- | 9-1 | Cattle | 2014 | 34 | + (*pheR*) | 40 kb | AMP, STR, SUL, TET | SAMD00234640/DRX226645 |
| L-4612 | 4,[5],12:i:- | 9-1 | Cattle | 2013 | 34 | + (*pheR*) | 40 kb | AMP, STR, SUL, TET | SAMD00234682/DRX226687 |
| L-4650 | 4,[5],12:i:- | 9-1 | Cattle | 2015 | 34 | + (*pheR*) | 40 kb | AMP, STR, SUL, TET | SAMD00234713/DRX226718 |
| L-4647 | 4,[5],12:i:- | 9-1 | Cattle | 2015 | 34 | + (*pheR*) | 40 kb | TET | SAMD00234710/DRX226715 |
| L-4353 | 4,[5],12:i:- | 9-1 | Cattle | 2015 | 34 | + (*pheR*) | 40 kb | AMP, STR, SUL, TET | SAMD00234585/DRX226590 |
| L-4351 | 4,[5],12:i:- | 9-1 | Cattle | 2015 | 34 | + (*pheR*) | 40 kb | AMP, STR, SUL | SAMD00234583/DRX226588 |
| L-4619 | 4,[5],12:i:- | 9-1 | Cattle | 2014 | 34 | + (*pheR*) | 40 kb | AMP, STR, SUL, TET | SAMD00234688/DRX226693 |
| L-4656 | 4,[5],12:i:- | 9-1 | Cattle | 2016 | 34 | + (*pheR*) | 40 kb | AMP, STR, SUL, TET | SAMD00234718/DRX226723 |
| L-4583 | 4,[5],12:i:- | 9-1 | Swine | 2016 | 34 | + (*pheR*) | 40 kb | AMP, STR, SUL, TET | SAMD00234658/DRX226663 |
| L-4688 | 4,[5],12:i:- | 9-1 | Cattle | 2017 | 34 | + (*pheR*) | 40 kb | AMP, STR, SUL, TET | SAMD00234743/DRX226748 |
| L-4697 | 4,[5],12:i:- | 9-1 | Cattle | 2017 | 34 | + (*pheR*) | 40 kb | AMP, SUL, TET | SAMD00234745/DRX226750 |
| L-4690 | 4,[5],12:i:- | 9-1 | Cattle | 2017 | 34 | + (*pheR*) | 40 kb | AMP, STR, SUL | SAMD00234744/DRX226749 |
| L-4633 | 4,[5],12:i:- | 9-1 | Cattle | 2015 | 34 | + (*pheR*) | 40 kb | AMP, STR, SUL, TET | SAMD00234699/DRX226704 |
| L-4664 | 4,[5],12:i:- | 9-1 | Cattle | 2016 | 34 | + (*pheR*) | 40 kb | AMP, STR, SUL, TET | SAMD00234725/DRX226730 |
| L-4362 | 4,[5],12:i:- | 9-1 | Cattle | 2015 | 34 | + (*pheR*) | 40 kb | AMP, STR, SUL, TET | SAMD00234592/DRX226597 |
| L-4366 | 4,[5],12:i:- | 9-1 | Cattle | 2016 | 34 | + (*pheR*) | 40 kb | AMP, STR, SUL, TET | SAMD00234594/DRX226599 |
| L-4361 | 4,[5],12:i:- | 9-1 | Cattle | 2015 | 34 | + (*pheR*) | 40 kb | AMP, STR, SUL, TET | SAMD00234591/DRX226596 |
| L-4634 | 4,[5],12:i:- | 9-1 | Cattle | 2015 | 34 | + (*pheR*) | 94 kb. 40 kb | AMP, STR, SUL, TET, CHL, TMP | SAMD00234700/DRX226705 |
| L-4649 | 4,[5],12:i:- | 9-1 | Cattle | 2015 | 34 | + (*pheR*) | 94 kb. 40 kb | STR, SUL, TET, CHL, TMP | SAMD00234712/DRX226717 |
| L-4654 | 4,[5],12:i:- | 9-1 | Cattle | 2016 | 34 | + (*pheR*) | 94 kb. 40 kb | AMP, STR, SUL, TET, CHL, TMP | SAMD00234716/DRX226721 |
| L-4670 | 4,[5],12:i:- | 9-1 | Cattle | 2016 | 34 | + (*pheR*) | 94 kb. 40 kb | AMP, STR, SUL, TET, CHL, TMP | SAMD00234730/DRX226735 |
| L-4668 | 4,[5],12:i:- | 9-1 | Cattle | 2016 | 34 | + (*pheR*) | 94 kb. 40 kb | AMP, STR, SUL, TET, CHL, TMP | SAMD00234728/DRX226733 |

*Continued on following pages*

**Technical Appendix Table 2**-Continued

| Strain | Serotype | Clade | Origin | Year | Sequence Type | ICEmST (locus)* | Plasmid profile[†] | Antimicrobial resistance pattern[‡] | BioSample/Experiment[§] accession no. or reference |
|---|---|---|---|---|---|---|---|---|---|
| L-4630 | 4,[5],12:i:- | 9-1 | Cattle | 2015 | 34 | + (*pheR*) | 94 kb. 40 kb | AMP, STR, SUL, TET, CHL, TMP | SAMD00234698/DRX226703 |
| L-4629 | 4,[5],12:i:- | 9-1 | Cattle | 2015 | 34 | + (*pheR*) | 94 kb. 40 kb | AMP, STR, SUL, TET, CHL, TMP | SAMD00234697/DRX226702 |
| L-4635 | 4,[5],12:i:- | 9-1 | Cattle | 2015 | 34 | + (*pheR*) | 94 kb. 40 kb | AMP, STR, SUL, TET, CHL, TMP | SAMD00234701/DRX226706 |
| L-4237 | 4,[5],12:i:- | 9-2 | Swine | 2012 | 34 | + (*pheR*) | ND | AMP, STR, SUL, TET | SAMD00234573/DRX226578 |
| L-4264 | 4,[5],12:i:- | 9-2 | Cattle | 2012 | 34 | + (*pheR*) | ND | AMP, STR, SUL, TET | SAMD00234574/DRX226579 |
| L-4328 | 4,[5],12:i:- | 9-2 | Swine | 2012 | 34 | + (*pheR*) | <40 kb (2) | AMP, STR, SUL, TET | SAMD00234576/DRX226581 |
| L-4330 | 4,[5],12:i:- | 9-2 | Cattle | 2013 | 34 | + (*pheR*) | ND | AMP, STR, SUL, TET | SAMD00234577/DRX226582 |
| L-4332 | 4,[5],12:i:- | 9-2 | Cattle | 2014 | 34 | + (*pheR*) | ND | AMP, STR, SUL, TET | SAMD00234578/DRX226583 |
| L-4676 | 4,[5],12:i:- | 9-2 | Cattle | 2014 | 34 | + (*pheR*) | 176 kb | AMP, STR, SUL, TET | SAMD00234735/DRX226740 |
| L-4679 | 4,[5],12:i:- | 9-2 | Cattle | 2015 | 34 | + (*pheR*) | <40k (2) | AMP, STR, SUL, TET, NAL | SAMD00234737/DRX226742 |
| L-4336 | 4,[5],12:i:- | 9-2 | Cattle | 2015 | 34 | + (*pheR*) | ND | AMP, STR, SUL, TET | SAMD00234581/DRX226586 |
| L-4333 | 4,[5],12:i:- | 9-2 | Cattle | 2014 | 34 | + (*pheR*) | 62 kb | AMP, STR, SUL, TET | SAMD00234579/DRX226584 |
| L-4334 | 4,[5],12:i:- | 9-2 | Cattle | 2015 | 34 | + (*pheR*) | ND | AMP, SUL, TET | SAMD00234580/DRX226585 |
| L-4345 | 4,[5],12:i:- | 9-2 | Cattle | 2015 | 34 | + (*pheR*) | ND | AMP, STR, SUL, TET | SAMD00234582/DRX226587 |
| L-4683 | 4,[5],12:i:- | 9-2 | Swine | 2013 | 34 | + (*pheR*) | 176 kb | AMP, STR, SUL, TET | SAMD00234739/DRX226744 |
| L-4234 | 4,[5],12:i:- | 9-2 | Swine | 2014 | 34 | + (*pheR*) | ND | AMP, STR, SUL, TET | 19 |
| L-4471 | 4,[5],12:i:- | 9-2 | Cattle | 2016 | 34 | + (*pheR*) | <40 kb (1) | AMP, STR, SUL, TET | SAMD00234613/DRX226618 |
| L-4469 | 4,[5],12:i:- | 9-2 | Cattle | 2015 | 34 | + (*pheR*) | <40 kb (1) | AMP, STR, SUL, TET | SAMD00234612/DRX226617 |
| L-4266 | 4,[5],12:i:- | 9-2 | Cattle | 2015 | 34 | + (*pheR*) | 62 kb | AMP, STR, SUL, TET | SAMD00234575/DRX226580 |
| L-4473 | 4,[5],12:i:- | 9-2 | Swine | 2016 | 34 | + (*pheR*) | ND | SUL, TET | SAMD00234614/DRX226619 |
| L-4685 | 4,[5],12:i:- | 9-2 | Swine | 2016 | 34 | + (*pheR*) | <40 kb (2) | AMP, STR, SUL, TET | SAMD00234741/DRX226746 |
| L-4598 | 4,[5],12:i:- | 9-2 | Cattle | 2016 | 34 | + (*pheR*) | ND | SUL, TET | SAMD00234672/DRX226677 |
| L-4605 | 4,[5],12:i:- | 9-2 | Cattle | 2016 | 34 | + (*pheR*) | 94 kb | AMP, SUL, TET, GEN | SAMD00234679/DRX226684 |
| L-4565 | 4,[5],12:i:- | 9-2 | Cattle | 2014 | 34 | + (*pheR*) | ND | AMP, STR, SUL, TET | SAMD00234643/DRX226648 |
| L-4628 | 4,[5],12:i:- | 9-2 | Cattle | 2015 | 34 | - | ND | AMP, STR, SUL, TET | SAMD00234696/DRX226701 |
| L-4233 | 4,[5],12:i:- | 9-2 | Cattle | 2014 | 34 | + (*pheR*) | <40 kb (1) | AMP, STR, SUL, TET | 19 |
| L-4741 | 4,[5],12:i:- | 9-2 | Cattle | 2015 | 34 | + (*pheR*) | ND | AMP, SUL | SAMD00234746/DRX226751 |

*Continued on following pages*

**Technical Appendix Table 2**-Continued

| Strain | Serotype | Clade | Origin | Year | Sequence Type | ICEmST (locus)* | Plasmid profile † | Antimicrobial resistance pattern ‡ | BioSample/Experiment§ accession no. or reference |
|---|---|---|---|---|---|---|---|---|---|
| L-4795 | 4,[5],12:i:- | 9-2 | Swine | 2017 | 34 | + (*pheR*) | <40 kb (3) | AMP, STR, SUL, TET, CHL, TMP | SAMD00234764/DRX226769 |
| L-4792 | 4,[5],12:i:- | 9-2 | Swine | 2017 | 34 | + (*pheR*) | <40 kb (3) | AMP, STR, SUL, TET | SAMD00234761/DRX226766 |
| L-4794 | 4,[5],12:i:- | 9-2 | Swine | 2017 | 34 | + (*pheR*) | <40 kb (3) | AMP, STR, SUL, TET | SAMD00234763/DRX226768 |
| L-4784 | 4,[5],12:i:- | 9-2 | Swine | 2017 | 34 | + (*pheR*) | <40 kb (3) | AMP, STR, SUL, TET | SAMD00234753/DRX226758 |
| L-4793 | 4,[5],12:i:- | 9-2 | Swine | 2017 | 34 | + (*pheR*) | <40 kb (3) | AMP, STR, SUL, TET | SAMD00234762/DRX226767 |
| L-4782 | 4,[5],12:i:- | 9-2 | Swine | 2017 | 34 | + (*pheR*) | <40 kb (3) | TET | SAMD00234751/DRX226756 |
| L-4785 | 4,[5],12:i:- | 9-2 | Swine | 2017 | 34 | + (*pheR*) | <40 kb (3) | Susceptible | SAMD00234754/DRX226759 |
| L-4786 | 4,[5],12:i:- | 9-2 | Swine | 2017 | 34 | + (*pheR*) | <40 kb (3) | TET | SAMD00234755/DRX226760 |
| L-4542 | 4,[5],12:i:- | 9-2 | Swine | 2016 | 34 | + (*pheR*) | <40 kb (3) | SUL, TET | SAMD00234623/DRX226628 |
| L-4545 | 4,[5],12:i:- | 9-2 | Swine | 2016 | 34 | + (*pheR*) | <40 kb (3) | SUL | SAMD00234624/DRX226629 |
| L-4788 | 4,[5],12:i:- | 9-2 | Swine | 2017 | 34 | + (*pheR*) | <40 kb (3) | AMP, STR, SUL, TET | SAMD00234757/DRX226762 |
| L-4787 | 4,[5],12:i:- | 9-2 | Swine | 2017 | 34 | + (*pheR*) | <40 kb (3) | AMP, STR, SUL, TET | SAMD00234756/DRX226761 |
| L-4777 | 4,[5],12:i:- | 9-2 | Swine | 2017 | 34 | + (*pheR*) | <40 kb (4) | AMP, STR, SUL, TET, CHL, TMP | SAMD00234747/DRX226752 |
| L-4538 | 4,[5],12:i:- | 9-2 | Swine | 2017 | 34 | + (*pheR*) | <40 kb (3) | AMP, STR, SUL, TET | SAMD00234620/DRX226625 |
| L-4791 | 4,[5],12:i:- | 9-2 | Swine | 2017 | 34 | + (*pheR*) | ND | AMP, STR, SUL, TET | SAMD00234760/DRX226765 |
| L-4790 | 4,[5],12:i:- | 9-2 | Swine | 2017 | 34 | + (*pheR*) | ND | AMP, STR, SUL, TET | SAMD00234759/DRX226764 |
| L-4789 | 4,[5],12:i:- | 9-2 | Swine | 2017 | 34 | + (*pheR*) | ND | AMP, STR, SUL, TET | SAMD00234758/DRX226763 |
| L-4539 | 4,[5],12:i:- | 9-2 | Swine | 2016 | 34 | + (*pheR*) | <40 kb (3) | AMP, STR, SUL, TET | SAMD00234621/DRX226626 |
| L-4783 | 4,[5],12:i:- | 9-2 | Swine | 2017 | 34 | + (*pheR*) | 94 kb, <40 kb (3) | AMP, STR, SUL, TET | SAMD00234752/DRX226757 |
| L-4540 | 4,[5],12:i:- | 9-2 | Swine | 2016 | 34 | + (*pheR*) | <40 kb (2) | AMP, STR, SUL, TET | SAMD00234622/DRX226627 |
| L-4536 | 4,[5],12:i:- | 9-2 | Swine | 2016 | 34 | + (*pheR*) | <40 kb (3) | AMP, STR, SUL, TET | SAMD00234619/DRX226624 |
| L-4780 | 4,[5],12:i:- | 9-2 | Swine | 2017 | 34 | + (*pheR*) | <40 kb (4) | AMP, STR, SUL, TET, CHL, TMP | SAMD00234749/DRX226754 |
| L-4781 | 4,[5],12:i:- | 9-2 | Swine | 2017 | 34 | + (*pheR*) | <40 kb (4) | AMP, STR, SUL, TET, CHL, TMP | SAMD00234750/DRX226755 |
| L-4779 | 4,[5],12:i:- | 9-2 | Swine | 2017 | 34 | + (*pheR*) | <40 kb (4) | AMP, STR, SUL, TET, CHL, TMP | SAMD00234748/DRX226753 |
| L-4548 | 4,[5],12:i:- | 9-2 | Cattle | 2012 | 34 | + (*pheR*) | ND | AMP, STR, SUL, TET | SAMD00234627/DRX226632 |
| L-4440 | 4,[5],12:i:- | 9-2 | Cattle | 2012 | 34 | + (*pheR*) | ND | AMP, STR, SUL, TET | SAMD00234598/DRX226603 |
| L-4475 | 4,[5],12:i:- | 9-2 | Cattle | 2012 | 34 | + (*pheR*) | ND | AMP, STR, SUL, TET | SAMD00234615/DRX226620 |

*Continued on following pages*

**Technical Appendix Table 2**-Continued

| Strain | Serotype | Clade | Origin | Year | Sequence Type | ICEmST (locus)* | Plasmid profile † | Antimicrobial resistance pattern ‡ | BioSample/Experiment§ accession no. or reference |
|---|---|---|---|---|---|---|---|---|---|
| L-4467 | 4,[5],12:i:- | 9-2 | Cattle | 2014 | 34 | + (*pheR*) | ND | AMP, STR, SUL, TET | SAMD00234611/DRX226616 |
| L-4465 | 4,[5],12:i:- | 9-2 | Cattle | 2013 | 34 | + (*pheR*) | ND | AMP, STR, SUL, TET | SAMD00234610/DRX226615 |
| L-4259 | 4,[5],12:i:- | 9-2 | Cattle | 2013 | 34 | + (*pheR*) | ND | AMP, STR, SUL, TET | 19 |
| L-4236 | 4,[5],12:i:- | 9-2 | Swine | 2013 | 34 | + (*pheR*) | ND | AMP, STR, SUL, TET | SAMD00234572/DRX226577 |
| L-4261 | 4,[5],12:i:- | 9-2 | Cattle | 2014 | 34 | + (*pheR*) | 62 kb, <40 kb (2) | AMP, STR, SUL, TET, KAN | 19 |
| L-4257 | 4,[5],12:i:- | 9-2 | Cattle | 2013 | 34 | + (*pheR*) | ND | AMP, STR, SUL, TET | 19 |
| L-4561 | 4,[5],12:i:- | 9-2 | Swine | 2014 | 34 | + (*pheR*) | ND | AMP, STR, SUL, TET | SAMD00234639/DRX226644 |
| L-4443 | 4,[5],12:i:- | 9-2 | Swine | 2014 | 34 | + (*pheR*) | ND | AMP, STR, SUL, TET | SAMD00234601/DRX226606 |
| L-4447 | 4,[5],12:i:- | 9-2 | Swine | 2015 | 34 | + (*pheR*) | ND | AMP, STR, SUL, TET | SAMD00234605/DRX226610 |
| L-4572 | 4,[5],12:i:- | 9-2 | Swine | 2015 | 34 | + (*pheR*) | ND | AMP, STR, SUL, TET | SAMD00234650/DRX226655 |
| L-4601 | 4,[5],12:i:- | 9-2 | Swine | 2016 | 34 | + (*pheV*) | <40 kb (1) | SUL, TET | SAMD00234675/DRX226680 |
| L-4585 | 4,[5],12:i:- | 9-2 | Swine | 2016 | 34 | + (*pheR*) | ND | AMP, STR, SUL, TET | SAMD00234660/DRX226665 |
| L-4558 | 4,[5],12:i:- | 9-2 | Swine | 2013 | 34 | + (*pheR*) | ND | AMP, STR, SUL, TET | SAMD00234636/DRX226641 |
| L-4555 | 4,[5],12:i:- | 9-2 | Swine | 2013 | 34 | + (*pheR*) | ND | AMP, STR, SUL, TET | SAMD00234633/DRX226638 |
| L-4559 | 4,[5],12:i:- | 9-2 | Cattle | 2014 | 34 | + (*pheR*) | ND | AMP, STR, SUL, TET | SAMD00234637/DRX226642 |
| L-4560 | 4,[5],12:i:- | 9-2 | Cattle | 2014 | 34 | + (*pheR*) | ND | AMP, STR, SUL, TET | SAMD00234638/DRX226643 |
| L-4564 | 4,[5],12:i:- | 9-2 | Cattle | 2014 | 34 | + (*pheR*) | ND | AMP, STR, SUL, TET | SAMD00234642/DRX226647 |
| L-4569 | 4,[5],12:i:- | 9-2 | Cattle | 2015 | 34 | + (*pheR*) | ND | AMP, STR, SUL, TET | SAMD00234647/DRX226652 |
| L-4446 | 4,[5],12:i:- | 9-2 | Cattle | 2015 | 34 | + (*pheR*) | ND | AMP, STR, SUL, TET | SAMD00234604/DRX226609 |
| L-4570 | 4,[5],12:i:- | 9-2 | Cattle | 2015 | 34 | + (*pheR*) | ND | AMP, STR, SUL, TET | SAMD00234648/DRX226653 |
| L-4596 | 4,[5],12:i:- | 9-2 | Cattle | 2016 | 34 | + (*pheR*) | 208 kb | AMP, STR, SUL, TET, CHL, FOF | SAMD00234670/DRX226675 |
| L-4586 | 4,[5],12:i:- | 9-2 | Cattle | 2016 | 34 | + (*pheR*) | ND | AMP, STR, SUL, TET | SAMD00234661/DRX226666 |
| L-4450 | 4,[5],12:i:- | 9-2 | Cattle | 2016 | 34 | + (*pheR*) | ND | AMP, STR, SUL, TET | SAMD00234608/DRX226613 |
| L-4587 | 4,[5],12:i:- | 9-2 | Cattle | 2016 | 34 | + (*pheR*) | ND | AMP, STR, SUL, TET | SAMD00234662/DRX226667 |
| L-4445 | 4,[5],12:i:- | 9-2 | Cattle | 2014 | 34 | + (*pheR*) | 176 kb, <40 kb (1) | AMP, STR, SUL, TET, CHL, TMP | SAMD00234603/DRX226608 |
| L-4567 | 4,[5],12:i:- | 9-2 | Cattle | 2014 | 34 | + (*pheR*) | 176 kb, <40 kb (1) | AMP, STR, SUL, TET, CHL, TMP, GEN | SAMD00234645/DRX226650 |
| L-4620 | 4,[5],12:i:- | 9-2 | Cattle | 2014 | 34 | + (*pheR*) | ND | AMP, STR, SUL, TET | SAMD00234689/DRX226694 |

*Continued on following pages*

**Technical Appendix Table 2**-Continued

| Strain | Serotype | Clade | Origin | Year | Sequence Type | ICEmST (locus)* | Plasmid profile † | Antimicrobial resistance pattern ‡ | BioSample/Experiment§ accession no. or reference |
|---|---|---|---|---|---|---|---|---|---|
| L-4357 | 4,[5],12:i:- | 9-2 | Cattle | 2015 | 34 | + (*pheR*) | ND | AMP, STR, SUL, TET | SAMD00234588/DRX226593 |
| L-4359 | 4,[5],12:i:- | 9-2 | Cattle | 2015 | 34 | + (*pheR*) | ND | AMP, STR, SUL, TET | SAMD00234589/DRX226594 |
| L-4652 | 4,[5],12:i:- | 9-2 | Cattle | 2015 | 34 | + (*pheR*) | ND | AMP, STR, SUL, TET | SAMD00234714/DRX226719 |
| L-4653 | 4,[5],12:i:- | 9-2 | Cattle | 2016 | 34 | + (*pheR*) | ND | AMP, STR, SUL, TET | SAMD00234715/DRX226720 |
| L-4655 | 4,[5],12:i:- | 9-2 | Cattle | 2016 | 34 | + (*pheR*) | ND | AMP, STR, SUL, TET | SAMD00234717/DRX226722 |
| L-4352 | 4,[5],12:i:- | 9-2 | Cattle | 2015 | 34 | + (*pheR*) | ND | AMP, STR, SUL, TET | SAMD00234584/DRX226589 |
| L-4354 | 4,[5],12:i:- | 9-2 | Cattle | 2015 | 34 | + (*pheR*) | ND | AMP, STR, SUL, TET | SAMD00234586/DRX226591 |
| L-4574 | 4,[5],12:i:- | 9-2 | Cattle | 2015 | 34 | + (*pheR*) | ND | AMP, STR, SUL, TET | SAMD00234652/DRX226657 |
| L-4448 | 4,[5],12:i:- | 9-2 | Cattle | 2015 | 34 | + (*pheR*) | ND | AMP, STR, SUL, TET | SAMD00234606/DRX226611 |
| L-4573 | 4,[5],12:i:- | 9-2 | Cattle | 2015 | 34 | + (*pheR*) | ND | AMP, STR, SUL, TET | SAMD00234651/DRX226656 |
| L-4580 | 4,[5],12:i:- | 9-2 | Cattle | 2015 | 34 | + (*pheR*) | ND | AMP, STR, SUL, TET | SAMD00234656/DRX226661 |
| L-4449 | 4,[5],12:i:- | 9-2 | Swine | 2015 | 34 | + (*pheR*) | ND | AMP, STR, SUL, TET | SAMD00234607/DRX226612 |
| L-4575 | 4,[5],12:i:- | 9-2 | Cattle | 2015 | 34 | + (*pheR*) | ND | AMP, STR, SUL, TET | SAMD00234653/DRX226658 |
| L-4576 | 4,[5],12:i:- | 9-2 | Swine | 2015 | 34 | + (*pheR*) | ND | AMP, STR, SUL, TET | SAMD00234654/DRX226659 |
| L-4595 | 4,[5],12:i:- | 9-2 | Cattle | 2016 | 34 | + (*pheR*) | ND | AMP, STR, SUL, TET | SAMD00234669/DRX226674 |
| L-4452 | 4,[5],12:i:- | 9-2 | Cattle | 2016 | 34 | + (*pheR*) | ND | AMP, STR, SUL, TET | SAMD00234609/DRX226614 |
| L-4666 | 4,[5],12:i:- | 9-2 | Cattle | 2016 | 34 | + (*pheR*) | ND | AMP, STR, SUL, TET | SAMD00234726/DRX226731 |
| L-4591 | 4,[5],12:i:- | 9-2 | Cattle | 2016 | 34 | + (*pheR*) | ND | AMP, STR, SUL, TET | SAMD00234665/DRX226670 |
| L-4593 | 4,[5],12:i:- | 9-2 | Cattle | 2016 | 34 | + (*pheR*) | ND | AMP, STR, SUL, TET | SAMD00234667/DRX226672 |
| L-4589 | 4,[5],12:i:- | 9-2 | Cattle | 2016 | 34 | + (*pheR*) | ND | AMP, STR, SUL, TET | SAMD00234664/DRX226669 |
| L-4441 | 4,[5],12:i:- | 9-2 | Cattle | 2012 | 34 | + (*pheR*) | ND | AMP, STR, SUL, TET | SAMD00234599/DRX226604 |
| L-4550 | 4,[5],12:i:- | 9-2 | Cattle | 2013 | 34 | + (*pheR*) | ND | AMP, STR, SUL, TET | SAMD00234629/DRX226634 |
| L-4549 | 4,[5],12:i:- | 9-2 | Cattle | 2012 | 34 | + (*pheR*) | ND | AMP, STR, SUL, TET | SAMD00234628/DRX226633 |
| L-4442 | 4,[5],12:i:- | 9-2 | Cattle | 2013 | 34 | + (*pheR*) | ND | AMP, STR, SUL, TET | SAMD00234600/DRX226605 |
| L-4552 | 4,[5],12:i:- | 9-2 | Cattle | 2013 | 34 | + (*pheR*) | ND | AMP, STR, SUL, TET | SAMD00234631/DRX226636 |
| L-4553 | 4,[5],12:i:- | 9-2 | Cattle | 2013 | 34 | + (*pheR*) | ND | AMP, STR, SUL, TET | SAMD00234632/DRX226637 |
| L-4557 | 4,[5],12:i:- | 9-2 | Cattle | 2013 | 34 | + (*pheR*) | ND | AMP, STR, SUL, TET | SAMD00234635/DRX226640 |

*Continued on following pages*

**Technical Appendix Table 2**-Continued

| Strain | Serotype | Clade | Origin | Year | Sequence Type | ICEmST (locus)* | Plasmid profile[†] | Antimicrobial resistance pattern[‡] | BioSample/Experiment[§] accession no. or reference |
|---|---|---|---|---|---|---|---|---|---|
| L-4615 | 4,[5],12:i:- | 9-2 | Cattle | 2013 | 34 | + (*pheR*) | ND | AMP, STR, SUL, TET | SAMD00234685/DRX226690 |
| L-4610 | 4,[5],12:i:- | 9-2 | Cattle | 2013 | 34 | + (*pheR*) | ND | AMP, STR, SUL, TET | SAMD00234680/DRX226685 |
| L-4617 | 4,[5],12:i:- | 9-2 | Cattle | 2014 | 34 | + (*pheR*) | ND | TET | SAMD00234686/DRX226691 |
| L-4614 | 4,[5],12:i:- | 9-2 | Cattle | 2013 | 34 | + (*pheR*) | 176 kb | AMP, STR, SUL, TET, TMP, KAN | SAMD00234684/DRX226689 |
| L-4624 | 4,[5],12:i:- | 9-2 | Cattle | 2014 | 34 | + (*pheR*) | ND | AMP, STR, SUL, TET | SAMD00234692/DRX226697 |
| L-4625 | 4,[5],12:i:- | 9-2 | Cattle | 2014 | 34 | + (*pheR*) | ND | AMP, STR, SUL, TET | SAMD00234693/DRX226698 |
| L-4626 | 4,[5],12:i:- | 9-2 | Cattle | 2014 | 34 | + (*pheR*) | 176 kb | AMP, STR, SUL, TET, CHL, TMP, GEN | SAMD00234694/DRX226699 |
| L-4584 | 4,[5],12:i:- | 9-2 | Cattle | 2016 | 34 | + (*pheR*) | 176 kb | AMP, STR, SUL, TET, CHL, TMP, GEN | SAMD00234659/DRX226664 |
| L-4588 | 4,[5],12:i:- | 9-2 | Cattle | 2016 | 34 | + (*pheR*) | 176 kb | AMP, STR, SUL, TET, CHL, TMP | SAMD00234663/DRX226668 |
| L-4597 | 4,[5],12:i:- | 9-2 | Cattle | 2016 | 34 | + (*pheR*) | 176 kb | AMP, STR, SUL, TET, CHL, TMP, GEN | SAMD00234671/DRX226676 |
| L-4640 | 4,[5],12:i:- | 9-2 | Cattle | 2015 | 34 | + (*pheR*) | 94 kb | AMP, STR, SUL, TET, CHL, TMP | SAMD00234706/DRX226711 |
| L-4667 | 4,[5],12:i:- | 9-2 | Cattle | 2016 | 34 | + (*pheR*) | 94 kb | AMP, STR, SUL, TET, CHL, TMP | SAMD00234727/DRX226732 |
| L-4639 | 4,[5],12:i:- | 9-2 | Cattle | 2015 | 34 | + (*pheR*) | 94 kb | AMP, STR, SUL, TET, CHL, TMP | SAMD00234705/DRX226710 |
| L-4641 | 4,[5],12:i:- | 9-2 | Cattle | 2015 | 34 | + (*pheR*) | 94 kb | AMP, STR, SUL, CHL, TMP | SAMD00234707/DRX226712 |
| L-4638 | 4,[5],12:i:- | 9-2 | Cattle | 2015 | 34 | + (*pheR*) | 94 kb | AMP, STR, SUL, TET, CHL, TMP | SAMD00234704/DRX226709 |
| L-4636 | 4,[5],12:i:- | 9-2 | Cattle | 2015 | 34 | + (*pheV*) | 94 kb | AMP, STR, SUL, TET, CHL, TMP | SAMD00234702/DRX226707 |
| L-4643 | 4,[5],12:i:- | 9-2 | Cattle | 2015 | 34 | + (*pheR*) | 94 kb | AMP, STR, SUL, TET, CHL, TMP | SAMD00234708/DRX226713 |
| L-4674 | 4,[5],12:i:- | 9-2 | Cattle | 2016 | 34 | + (*pheR*) | 94 kb, <40 kb (3) | AMP, STR, SUL, TET, CHL, TMP | SAMD00234733/DRX226738 |
| L-4592 | 4,[5],12:i:- | 9-2 | Cattle | 2016 | 34 | + (*pheR*) | 176 kb | AMP, STR, SUL, TET, CHL, TMP | SAMD00234666/DRX226671 |
| L-4599 | 4,[5],12:i:- | 9-2 | Cattle | 2016 | 34 | + (*pheR*) | 94 kb | AMP, STR, SUL, TET, CHL, TMP | SAMD00234673/DRX226678 |
| L-4600 | 4,[5],12:i:- | 9-2 | Cattle | 2016 | 34 | + (*pheR*) | 94 kb | AMP, STR, SUL, TET, CHL, TMP | SAMD00234674/DRX226679 |
| L-4603 | 4,[5],12:i:- | 9-2 | Cattle | 2016 | 34 | + (*pheR*) | 94 kb, 62 kb | AMP, STR, SUL, TET, CHL, TMP | SAMD00234677/DRX226682 |
| L-4675 | 4,[5],12:i:- | 9-2 | Cattle | 2016 | 34 | + (*pheR*) | 94 kb | AMP, STR, SUL, TET, CHL, TMP | SAMD00234734/DRX226739 |
| L-4657 | 4,[5],12:i:- | 9-2 | Cattle | 2016 | 34 | + (*pheR*) | 94 kb | AMP, STR, SUL, TET, CHL, TMP | SAMD00234719/DRX226724 |
| L-4669 | 4,[5],12:i:- | 9-2 | Cattle | 2016 | 34 | + (*pheR*) | 94 kb | AMP, STR, SUL, TET, CHL, TMP | SAMD00234729/DRX226734 |
| L-4662 | 4,[5],12:i:- | 9-2 | Cattle | 2016 | 34 | + (*pheR*) | 94 kb | AMP, STR, SUL, TET, CHL, TMP | SAMD00234723/DRX226728 |
| L-4661 | 4,[5],12:i:- | 9-2 | Cattle | 2016 | 34 | + (*pheR*) | 94 kb | AMP, STR, SUL, TET, CHL, TMP | SAMD00234722/DRX226727 |

*Continued on following pages*

**Technical Appendix Table 2**-Continued

| Strain | Serotype | Clade | Origin | Year | Sequence Type | ICEmST (locus)* | Plasmid profile † | Antimicrobial resistance pattern ‡ | BioSample/Experiment§ accession no. or reference |
|---|---|---|---|---|---|---|---|---|---|
| L-4660 | 4,[5],12:i:- | 9-2 | Cattle | 2016 | 34 | + (*pheR*) | 94 kb | AMP, STR, SUL, TET, CHL, TMP | SAMD00234721/DRX226726 |
| L-4627 | 4,[5],12:i:- | 9-2 | Cattle | 2015 | 34 | + (*pheR*) | ND | AMP, STR, SUL, TET | SAMD00234695/DRX226700 |
| L-4622 | 4,[5],12:i:- | 9-2 | Cattle | 2014 | 34 | + (*pheR*) | ND | AMP, STR, SUL, TET | SAMD00234691/DRX226696 |
| L-4568 | 4,[5],12:i:- | 9-2 | Cattle | 2015 | 34 | + (*pheR*) | ND | AMP, STR, SUL, TET | SAMD00234646/DRX226651 |
| L-4571 | 4,[5],12:i:- | 9-2 | Cattle | 2015 | 34 | + (*pheR*) | ND | AMP, STR, SUL, TET | SAMD00234649/DRX226654 |
| L-4582 | 4,[5],12:i:- | 9-2 | Swine | 2016 | 34 | + (*pheR*) | ND | AMP, STR, SUL, TET | SAMD00234657/DRX226662 |
| L-4578 | 4,[5],12:i:- | 9-2 | Cattle | 2015 | 34 | + (*pheR*) | 208 kb, <40 kb (1) | AMP, STR, SUL, TET, CHL, TMP, KAN | SAMD00234655/DRX226660 |
| L-4604 | 4,[5],12:i:- | 9-2 | Cattle | 2016 | 34 | + (*pheR*) | 208 kb | AMP, STR, SUL, TET, CHL, TMP, KAN | SAMD00234678/DRX226683 |
| L-4533 | 4,[5],12:i:- | 9-2 | Cattle | 2016 | 34 | + (*pheR*) | 208 kb | AMP, STR, SUL, TET, CHL, TMP, KAN | SAMD00234617/DRX226622 |
| L-4535 | 4,[5],12:i:- | 9-2 | Cattle | 2016 | 34 | + (*pheR*) | 208 kb | AMP, STR, SUL, TET, CHL, TMP, KAN | SAMD00234618/DRX226623 |
| L-4659 | 4,[5],12:i:- | 9-2 | Cattle | 2016 | 34 | + (*pheR*) | 208 kb | AMP, STR, SUL, TET, CHL, TMP, KAN | SAMD00234720/DRX226725 |
| L-4663 | 4,[5],12:i:- | 9-2 | Cattle | 2016 | 34 | + (*pheR*) | 208 kb | AMP, STR, SUL, TET, CHL, TMP, KAN | SAMD00234724/DRX226729 |
| L-4594 | 4,[5],12:i:- | 9-2 | Cattle | 2016 | 34 | + (*pheR*) | 208 kb | AMP, STR, SUL, TET, CHL, TMP, KAN | SAMD00234668/DRX226673 |
| L-4673 | 4,[5],12:i:- | 9-2 | Cattle | 2016 | 34 | + (*pheR*) | 94 kb | AMP, STR, SUL, TET, CHL, TMP, KAN | SAMD00234732/DRX226737 |
| L-4672 | 4,[5],12:i:- | 9-2 | Cattle | 2016 | 34 | + (*pheR*) | 94 kb (2) | AMP, STR, SUL, TET, CHL, TMP, KAN | SAMD00234731/DRX226736 |
| L-4618 | 4,[5],12:i:- | 9-2 | Cattle | 2014 | 34 | + (*pheR*) | ND | AMP, STR, SUL, TET | SAMD00234687/DRX226692 |
| L-4621 | 4,[5],12:i:- | 9-2 | Cattle | 2014 | 34 | + (*pheR*) | ND | AMP, STR, SUL, TET | SAMD00234690/DRX226695 |
| L-4646 | 4,[5],12:i:- | 9-2 | Cattle | 2015 | 34 | + (*pheR*) | ND | AMP, STR, SUL, TET | SAMD00234709/DRX226714 |
| L-4602 | 4,[5],12:i:- | 9-2 | Cattle | 2016 | 34 | + (*pheR*) | 94 kb | AMP, STR, SUL, TET, CHL, TMP | SAMD00234676/DRX226681 |
| L-4367 | 4,[5],12:i:- | 9-2 | Cattle | 2016 | 34 | + (*pheR*) | 94 kb | AMP, STR, SUL, TET, CHL, TMP | SAMD00234595/DRX226600 |
| L-4355 | 4,[5],12:i:- | 9-2 | Cattle | 2015 | 34 | + (*pheR*) | 94 kb | AMP, STR, SUL, TET | SAMD00234587/DRX226592 |
| L-4364 | 4,[5],12:i:- | 9-2 | Cattle | 2016 | 34 | + (*pheR*) | <40 kb (1) | AMP, STR, SUL, TET, CHL, TMP | SAMD00234593/DRX226598 |
| L-4360 | 4,[5],12:i:- | 9-2 | Cattle | 2015 | 34 | + (*pheR*) | 94 kb | AMP, STR, SUL, TET, CHL, TMP | SAMD00234590/DRX226595 |
| L-4648 | 4,[5],12:i:- | 9-2 | Cattle | 2015 | 34 | + (*pheR*) | 94 kb | AMP, STR, SUL, TET, CHL, TMP | SAMD00234711/DRX226716 |
| L-4637 | 4,[5],12:i:- | 9-2 | Cattle | 2015 | 34 | + (*pheR*) | 94 kb | AMP, STR, SUL, TET | SAMD00234703/DRX226708 |

*+, presence; -, absence; locus, location of ICEmST on the chromosome

†208 kb, 176 kb, 94 kb, 62 kb, 40 kb indicate the presence of each size plasmid; the number in parentheses indicate the number of plasmid; ND, not detected.

‡AMP, ampicillin; CFZ, cefazolin; CHL, chloramphenicol; CTX, cefotaxime; FEP, cefepime; GEN, gentamycin; KAN, kanamycin; NAL, nalidixic acid; STR, streptomycin;

SUL, sulfonamides; TET, tetracycline; TMP, trimethoprim

[§]The data have been deposited with links to DDBJ Sequence Read Archives accession number DRA010462.

**Technical Appendix Table 3.** Transfer frequency of ICEmST to *Salmonella* Typhimurium LT2 strain

| Donor strain | Clade | Average (per donor) | SD |
|---|---|---|---|
| L-3841 | 9-1 | $5.3\times10^{-7}$ | $5.7\times10^{-7}$ |
| L-3844 | 9-1 | $1.5\times10^{-7}$ | $1.4\times10^{-7}$ |
| L-4444 | 9-1 | $1.5\times10^{-6}$ | $2.5\times10^{-7}$ |
| L-4448 | 9-2 | $4.7\times10^{-7}$ | $3.0\times10^{-7}$ |
| L-4675 | 9-2 | $8.8\times10^{-7}$ | $1.1\times10^{-6}$ |
| L-4782 | 9-2 | $8.8\times10^{-7}$ | $6.2\times10^{-7}$ |

**Technical Appendix Table 4.** Characteristics of plasmids identified in *Salmonella* Typhimurium and *Salmonella* 4,[5],12:i:- clade 9 strains

| Plasmid name | Clade of host strain | Size (bp) | Antimicrobial resistance genes* | Replicon type |
|---|---|---|---|---|
| pSAL4551-1 | 9-1 | 128,709 | *aac*(6')-*Ib-cr*, *arr-3*, $bla_{OXA-1}$, *catB3*, *sul1* | IncHI2, IncHI2A |
| pSAL4681-1 | 9-1 | 151,734 | *aac*(3)-*IV*, *aac*(6')-*Ib-cr*, *aph*(3')-*Ia*, *aph*(4)-*Ia*, *arr-3*, $bla_{OXA-1}$,*catB3*, *dfrA12*, *sul1*, *sul2*, *sul3* | IncHI2, IncHI2A |
| pSAL4233-1 | 9-2 | 7,618 | ND* | IncQ1 |
| pSAL4445-1 | 9-2 | 131,843 | *aac*(3)-*IId*, *aadA1*, *aadA2*, $bla_{TEM1-B}$, *cmlA1*, *dfrA12*, *floR*, *mcr-3.1*, *sul2*, *sul3*, *tet*(A), *tet*(M) | IncFIB(AP001918) IncFIC(FII) |
| pSAL4261-1 | 9-2 | 60,222 | ND | IncFII(pCoo) |
| pSAL4261-2 | 9-2 | 6,835 | ND | ColRNAI |
| pSAL4261-4 | 9-2 | 4,060 | ND | Col8282 |
| pSAL4567-1 | 9-2 | 131,843 | *aac*(3)-*IId*, *aadA1*, *aadA2*, $bla_{TEM1-B}$, *cmlA1*, *dfrA12*, *floR*, *mcr-3.1*, *sul2*, *sul3*, *tet*(A), *tet*(M) | IncFIB(AP001918) IncFIC(FII) |
| pSAL4578-1 | 9-2 | 229,829 | *aadA2*, *aph*(3')-*Ia*, *aph*(3")-*Ib*, *aph*(6)-*Id*, $bla_{TEM1-B}$, *dfrA12*, *floR*, *sul1*, *sul2*, *tet*(A) | IncHI2, IncHI2A |
| pSAL4596-1 | 9-2 | 230,500 | *aadA1*, *catA1*, *cmlA1*, *mcr-1.1*, *mcr-5.1*, *mef*(B), *sul3*, *tet*(B), *tet*(M) | IncFIA(HI1), IncHI1A, IncHI1B(R27) |
| pSAL4605-1 | 9-2 | 99,132 | *aac*(3)-*IId*, $bla_{TEM1-B}$, *mcr-3.1*, *sul3* | IncFIB(AP001918) IncFIC(FII) |
| pSAL4614-1 | 9-2 | 229,829 | *aadA2*, *aph*(3')-*Ia*, *aph*(3")-*Ib*, *aph*(6)-*Id*, $bla_{TEM1-B}$, *dfrA12*, *floR*, *sul1*, *sul2*, *tet*(A) | IncHI2, IncHI2A |

*ND, not detected.

(a)

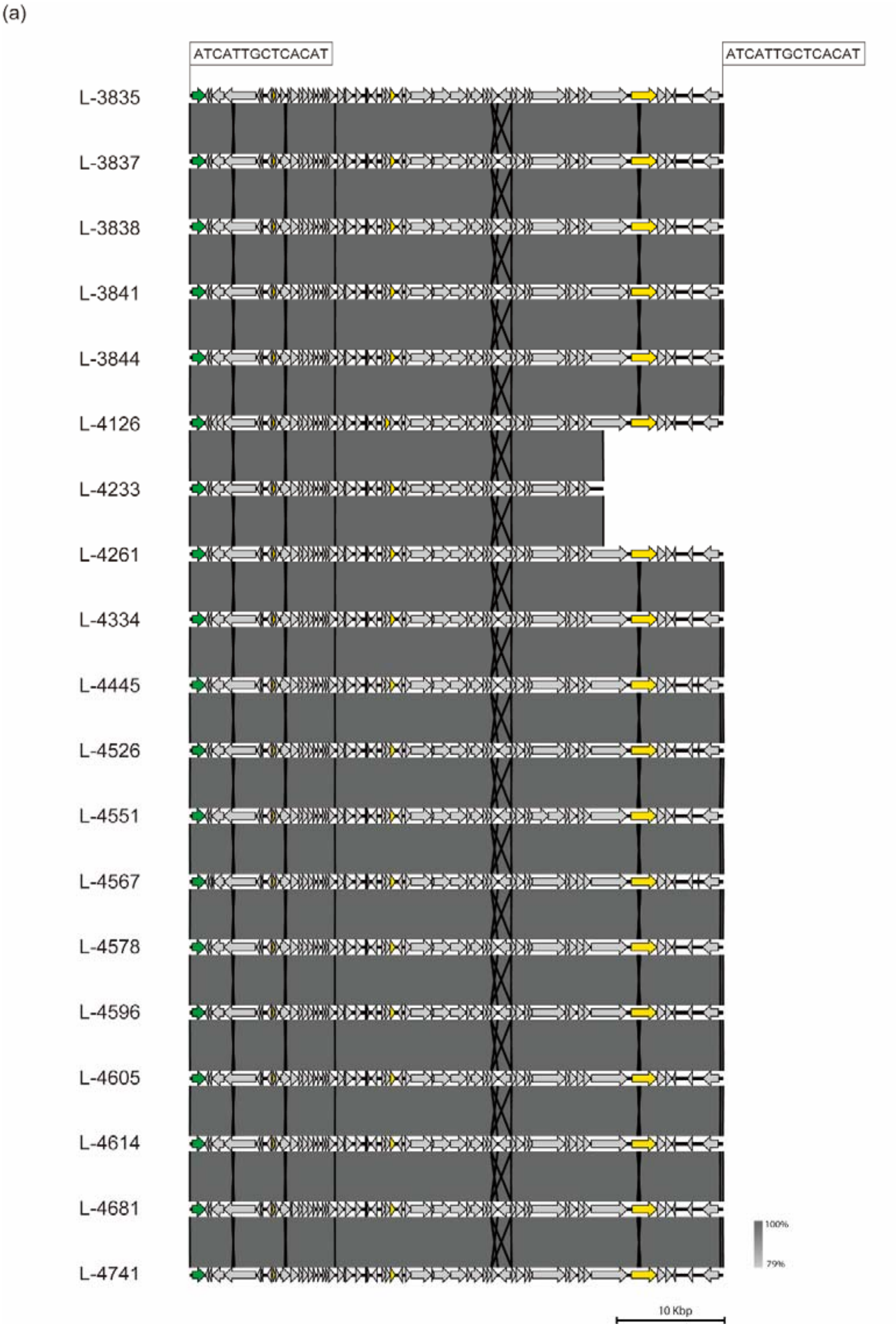


**Technical Appendix Figure 1**. Comparison of predicted prophages. The prophages which were identified as intact by PHASTER prophage searching tool were compared and visualized; (a) Gifsy-1, (b) Gifsy-2, (c) HP1, and (d) sal3. Green and gray arrows represent genes for integrase and the others, respectively. Yellow arrows indicate the ORFs which were used for homology search to investigate the prevalence of prophages in Figure 2. The nucleotide sequence shown in box are direct repeats which were probably generated upon integration of the prophage.

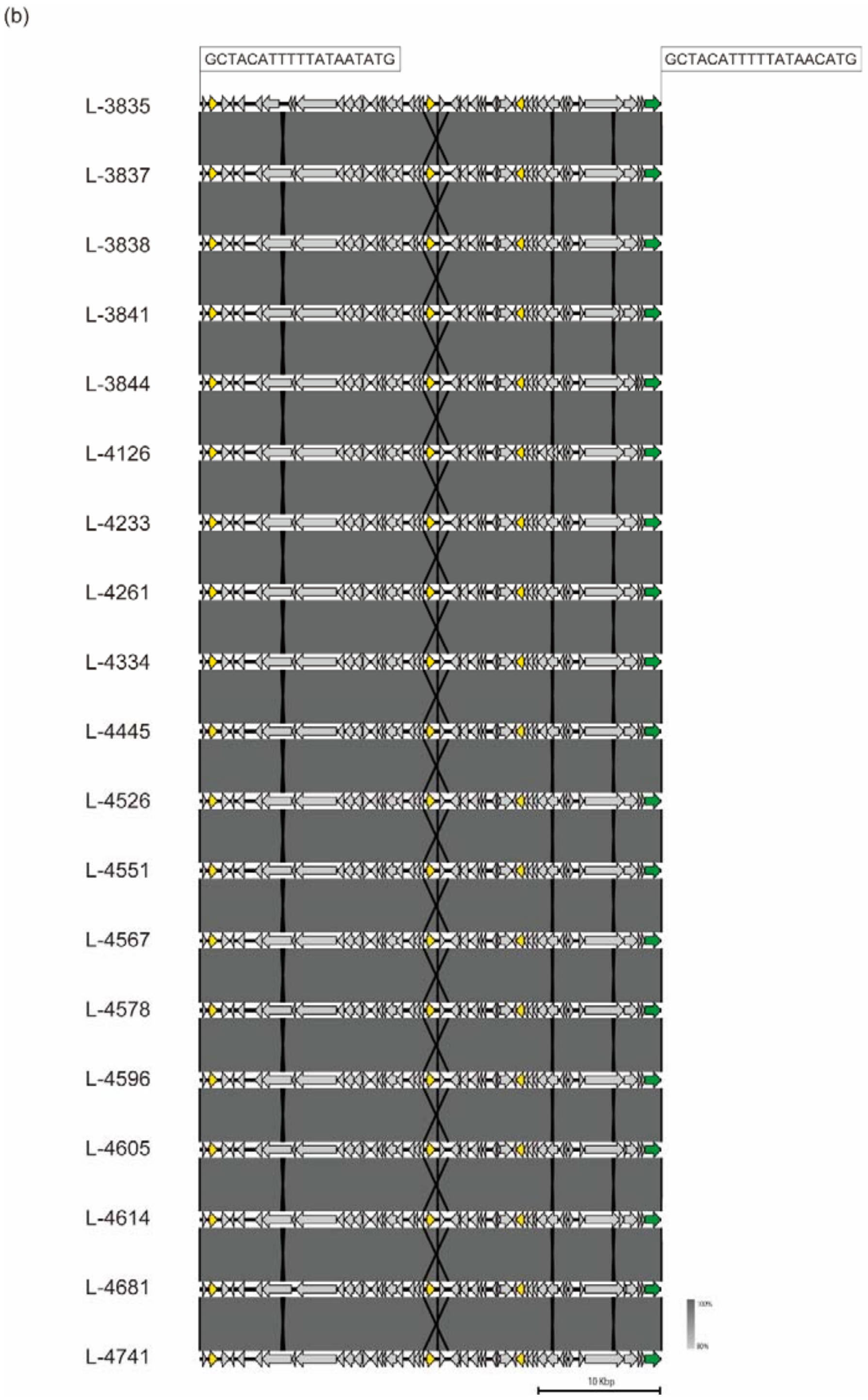


**Technical Appendix Figure 1**-Continued.

(c)

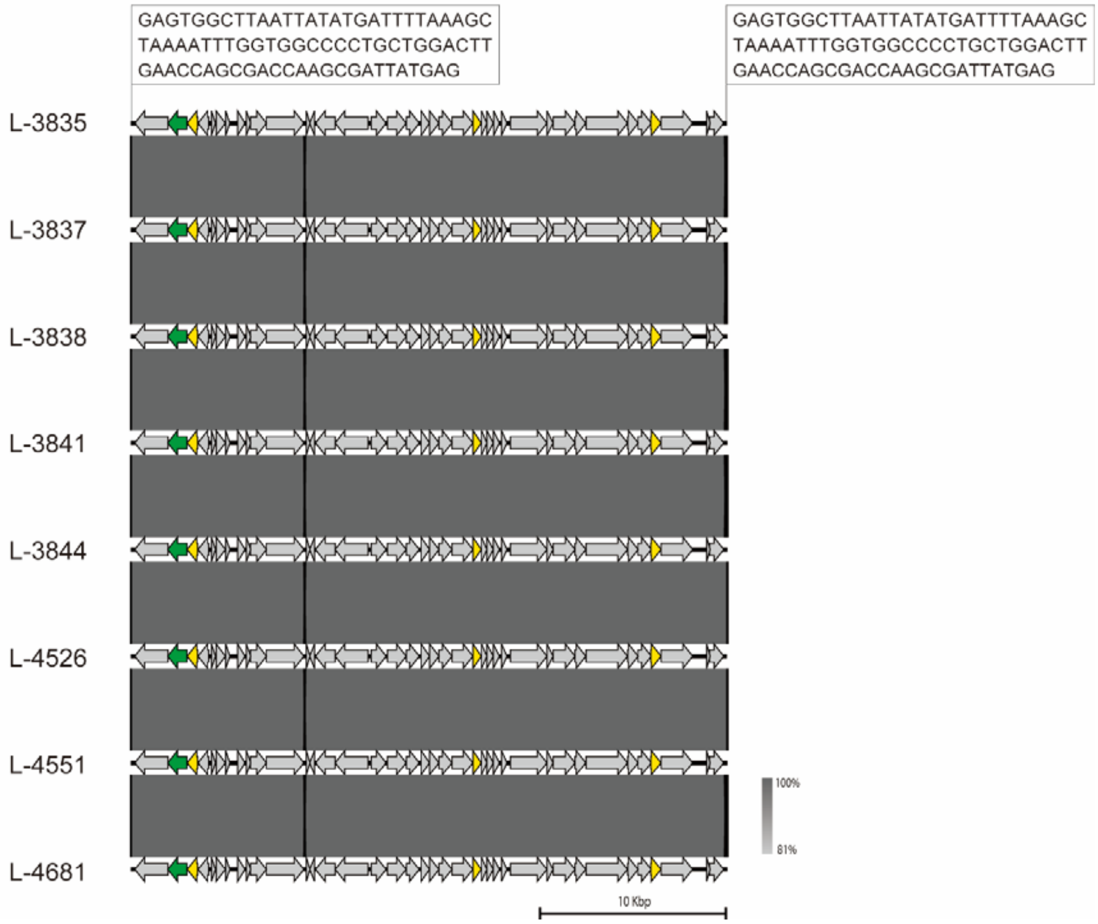


**Technical Appendix Figure 1**-Continued.

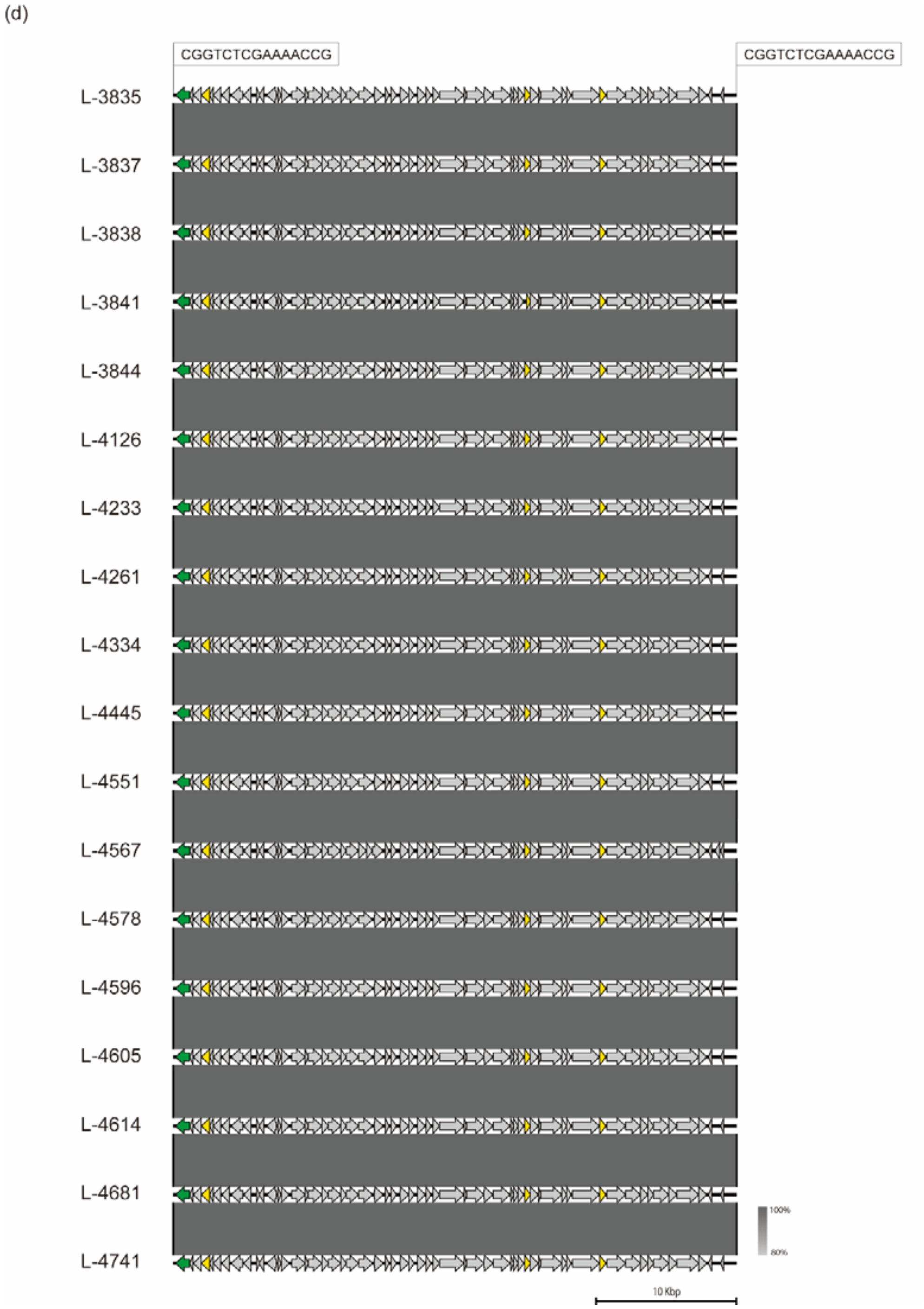


**Technical Appendix Figure 1**-Continued.
